# Early-Life Environmental Exposures Reprogram Epigenomic Aging to Alter Gene Expression Trajectories

**DOI:** 10.1101/2025.04.21.649824

**Authors:** Sandra L. Grimm, Rahul Jangid, Marisa S. Bartolomei, Dana C. Dolinoy, David L. Aylor, Gokhan M. Mutlu, Shyam Biswal, Bo Zhang, Ting Wang, TaRGET II Consortium, Cristian Coarfa, Cheryl Lyn Walker

## Abstract

To understand how early-life environmental exposures shape health and disease risk across the lifecourse, the TaRGET II Consortium exposed mice to diverse toxicants from pre-conception through weaning, and followed individual animals into adulthood, generating over 800 epigenomic and transcriptomic profiles. These profiles revealed that early-life exposures induced persistent epigenomic reprogramming and significantly disrupted the adult transcriptome. Notably, despite their diverse mechanisms of action, the exposure signatures of the xenoestrogen BPA, obesogen TBT, dioxin TCDD, and air pollutant PM2.5, were all largely comprised of genes normally differentially expressed during liver aging. Epigenetic histone modifications at enhancers—and, to a lesser extent, promoters—emerged as key targets for this reprogramming. Despite differing mechanisms of action, these four toxicants imparted similar “fingerprints” on the adult liver, characterized by direction-and cell type-specific polarization of the transcriptome. Hepatocyte genes that typically increase with age, particularly those in metabolic pathways, were downregulated, while conversely, non-parenchymal cell genes that typically decrease with age, such as those involved in extracellular matrix production, were upregulated. A similar signature of anti-correlation with programmed aging aging was also found in the transcriptome of patients with liver disease and hepatocellular carcinoma (HCC), and was effective at distinguishing healthy from diseased human livers. These findings demonstrate that the plasticity of epigenomic aging is vulnerable to early-life environmental exposures, which can reprogram the epigenome with lasting impacts on the transcriptome, and disease risk, later in life.

## INTRODUCTION

Epigenetic programming, once established, is heritable, yet retains the intrinsic plasticity necessary for three fundamental biological processes: remodeling the epigenome for cell-type specification during development, resetting imprinted loci for parent-of-origin-specific allele expression, and programmed epigenomic aging that directs changes in gene expression across the life course ^1,2^. Beyond these essential biologies, the epigenome is also susceptible to reprogramming by environmental exposures, particularly during critical windows of development ^3^. Both human epidemiological and animal model studies have demonstrated that environmental factors—including nutritional status, stress, and toxicant exposure—can induce lasting changes in the epigenome, with effects that persist long after the exposure has ceased.

Targets for epigenetic reprogramming include DNA methylation, histone modifications, and non-coding RNAs. Importantly, perturbation of the epigenome can lock in abnormal gene expression patterns, contributing to disease susceptibility in later life ^4,5^. This persistent epigenetic reprogramming is implicated in a wide range of adverse health outcomes, including metabolic disorders, cardiovascular diseases, and cancer ^5,6^. In some instances, these changes may even be transmitted to subsequent generations, compounding the long-term effects of the initial exposure through transgenerational epigenetic inheritance ^7,8^.

The Toxicant Exposures and Responses by Genomic and Epigenomic Regulators of Transcription II (TaRGET II) consortium (T2C) was established by the National Institute of Environmental Health Sciences (NIEHS) to understand how the epigenome and transcriptome respond to early-life environmental exposures and evaluate epigenomic signatures as biomarkers of exposure and predictors of disease risk ^9^. TaRGET II investigated a wide array of environmental toxicants, with diverse mechanisms of action. These included ligand activation of nuclear hormone receptors—such as the estrogen receptor, peroxisome proliferator-activated receptor, and aryl hydrocarbon receptor— triggered by compounds bisphenol A (BPA), tributyltin (TBT), and dioxin (TCDD) ^10,11^. The consortium also explored toxicants inducing oxidative stress, such as particulate matter (PM2.5) ^12^, agents that disrupt normal enzymatic, ion channel, and receptor function, such as lead (Pb) ^13^, and the endocrine disrupting chemical (EDC) Di(2-ethylhexyl) phthalate (DEHP) ^14^. All exposures began pre-conception and continued through weaning, after which exposure ceased, and animals were followed longitudinally into early (5 months) and later (10 months) adulthood to assess lasting effects. Further details on the exposures and methodologies employed across the consortium can be found in the Materials and Methods and publicly accessible metadata through the TaRGET II data portal (https://dcc.targetepigenomics.org/) and an accompanying database (http://toxitarget.com/).

In mice, the liver undergoes distinct morphological and functional transitions during embryonic and perinatal development, with well-characterized transcriptional correlates for this period of liver maturation ^15^. Significant transcriptional changes also occur in very old animals-particularly between 12 to 24 months of life-as highlighted by recent comprehensive studies ^16,17^. However, less is known about the epigenomic and transcriptomic shifts occurring between weaning, early and later adulthood—the interval selected by the T2C to assess the persistent effects of early-life environmental exposures. Therefore to lay a foundation for the exposure studies, longitudinal profiling of normal livers was conducted from 3 weeks to 10 months, examining histone modifications (this study) and DNA methylation ^18^, as well as transcriptomic changes via RNA sequencing (RNA-seq). The eight exposure groups were two doses of BPA: 10 mg/kg-diet/day [BPA.hi] and 10 μg/kg-diet/day [BPA.lo], DEHP (5mg/kg-diet/day), Pb (32 ppm-drinking water), TBT (3 uM-drinking water), TCDD (1 ug/kg-diet/day), and PM2.5 exposures conducted by two different consortium sites—Johns Hopkins University (PM2.5-JHU, concentrated PM2.5: 150 µg/m^3^) and the University of Chicago (PM2.5-CHI, synthetic PM2.5: 150 µg/m^3^) (Supplemental Figure 1a). From these study animals, the Consortium profiled individual mice, and for the present analysis, generated 410 transcriptomes (RNA-seq for males and females) and 436 epigenomes (ChIP-seq of males) for the eight different exposure groups and matched vehicle controls, at 3 different ages (weaning at 3 weeks, early adulthood at 5 months and later adulthood at 10 months), all without pooling samples. This individualized deep-sequencing approach enabled detection of exposure-specific transcriptional and epigenomic reprogramming while preserving the ability to account for inter-individual biological variability.

## RESULTS

### Early-Life Exposures Cause Persistent Disruption of the Adult Liver Transcriptome

As the mouse liver ages from weaning (3 weeks) to early adulthood (5 months), many genes exhibit differential expression, as demonstrated by the principal component analysis (PCA) and heatmaps presented in Figure 1a, along with UpSet plots in Figure 1b, which show numbers of male-specific, female-specific and shared (i.e. occurring in both sexes) gene expression changes. In both sexes, the majority of differentially expressed genes (DEGs, FDR <0.05, fold change exceeding 1.5x) that changed as a function of age (agingDEGs) exhibited a decrease in expression during this period: 1,153 agingDEGs were significantly downregulated in young adulthood in both males and females, compared to 296 agingDEGs that increased significantly during this time (Figure 1b). Furthermore, the majority of agingDEGs exhibited sex-specificity: we found 799 male-specific genes decreased in expression versus 303 that increased, and 235 female-specific agingDEGs that decreased compared to 129 that increased. Table 1 highlights the top agingDEGs upregulated and downregulated in males and females, with the complete list of all DEGs available in the combined signature file in Supplemental Table 1.

**Figure 1.**
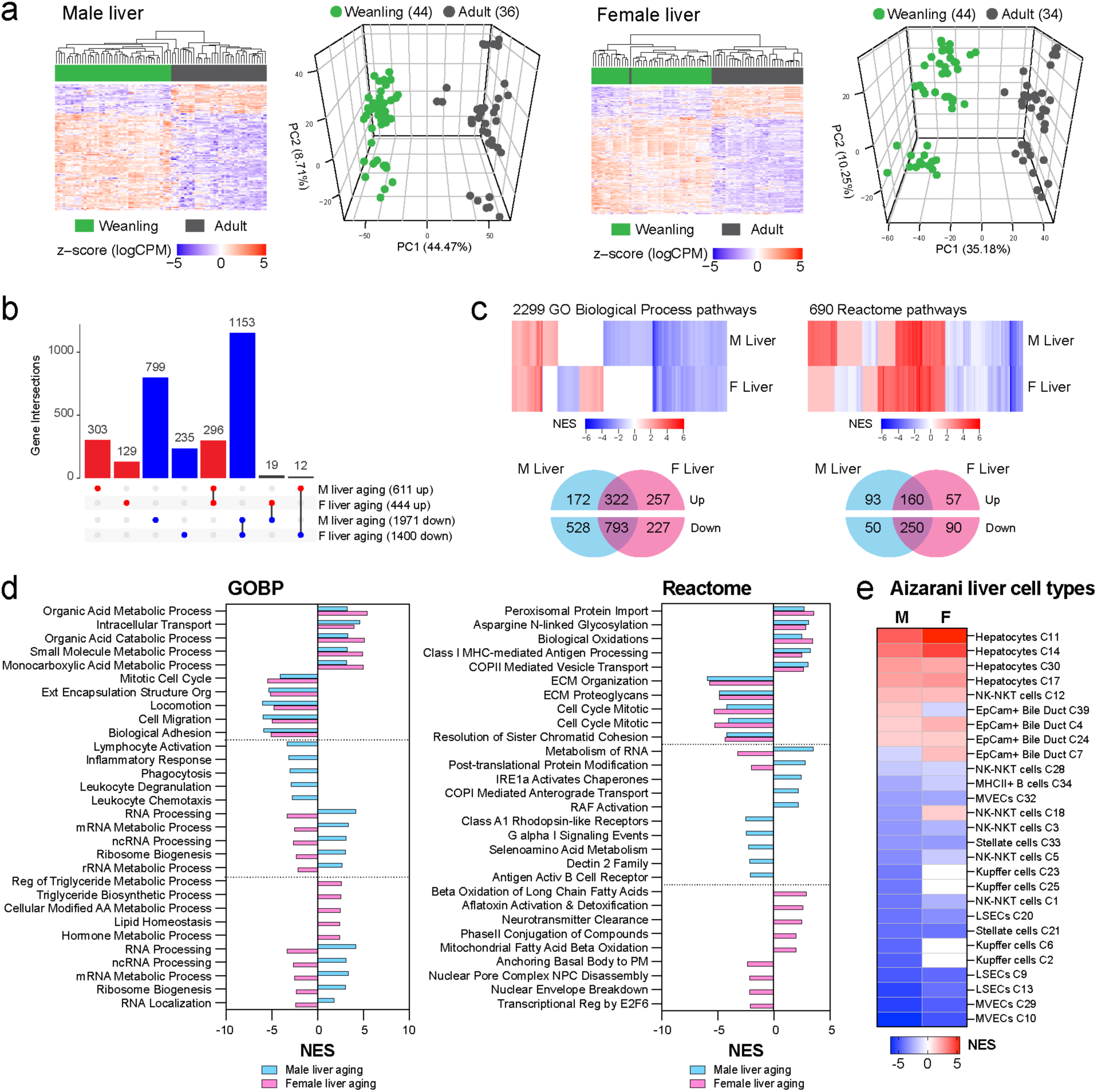
Transcriptomic signatures of aging in male and female mice in the 3-week to 5-month time window. a. PCA, heat maps and volcano plots of transcriptional changes associated with aging. b. UpSet plots comparing the aging transcriptome in males and females c. GSEA enriched pathways in male and female aging based on Gene Ontology Biological Processes (GOBP). d. Top shared and sex-specific GOBP and Reactome enriched pathways changed with age. e. GSEA enrichment results for liver cell type markers in male and female mice aging.

**Table 1.**
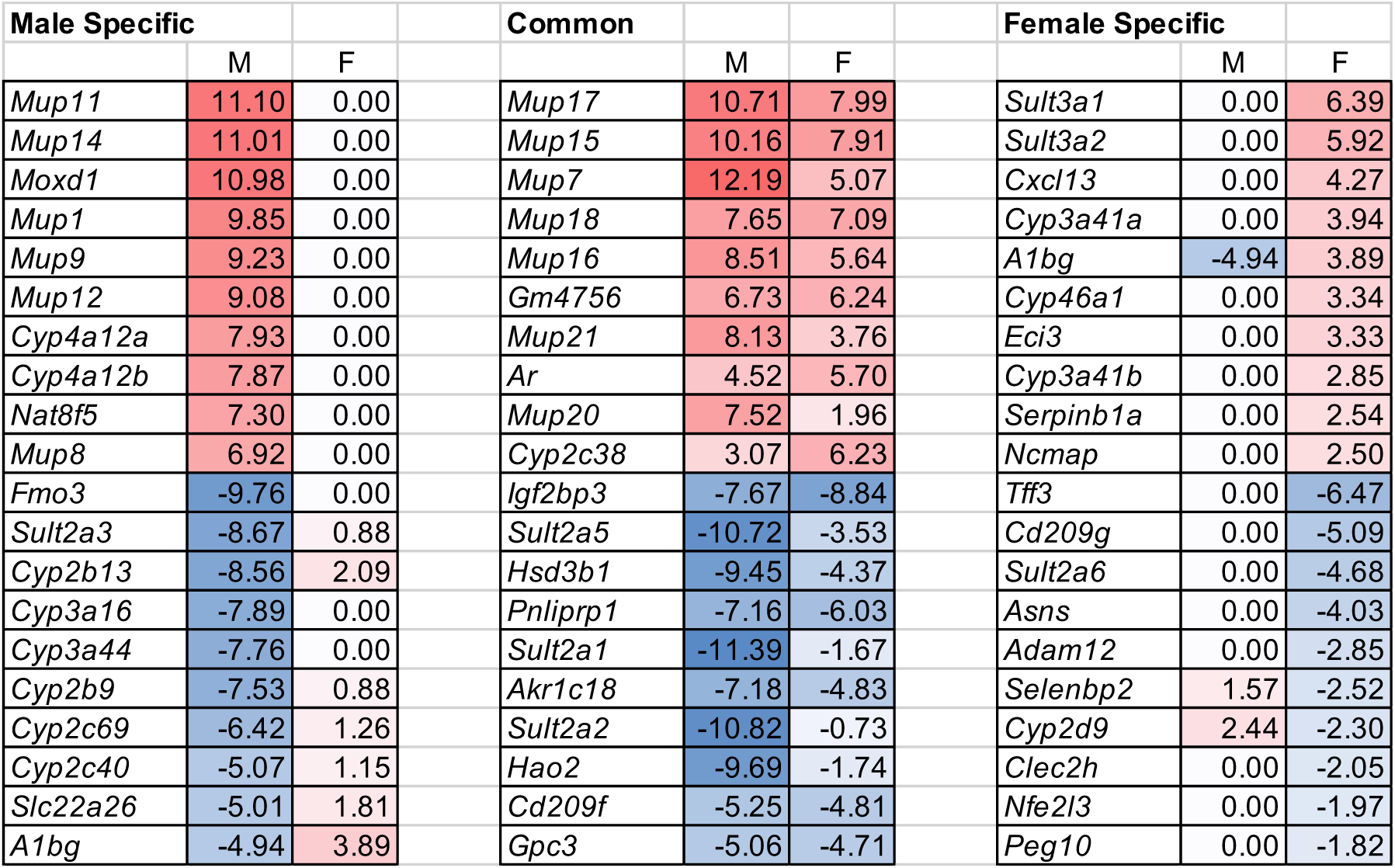
Top male and female liver aging DEGs.

Gene set enrichment analysis (GSEA) revealed many functional pathways that were positively or negatively enriched for these agingDEGs in both sexes. These are shown in Figure 1c-d, and Suppl Figure 1a-b; Supplemental Table 2 provides a complete list of enriched pathways and their core genes. Employing GO-BP and Reactome (Figure 1c-d) and Hallmark and KEGG compendia (Supplemental Figure 1b-c), pathways with the greatest enrichment in both males and females were associated with extracellular matrix production and cytoskeletal functions such as locomotion and mitosis (most negatively enriched) and cellular metabolism and peroxisome function (most positively enriched).

Interestingly, some pathways, such as those related to RNA processing, exhibited opposing trends in males (upregulated) versus females (downregulated) (Figure 1c). When GSEA was applied against cell identity genes for parenchymal (epithelial) and non-parenchymal liver cells, clear cell-type and directional specificity emerged. Between weaning and young adulthood, pathways enriched for parenchymal hepatocyte and bile duct epithelial cell identity genes were predominantly upregulated, whereas pathways enriched for cell identity genes of non-parenchymal cells, such as Stellate, Kupffer and endothelial cells, were downregulated (Figure 1e), consistent with the known metabolic, extracellular matrix production and immune signaling functions of these cell types.

We next assessed the impact of diverse early-life environmental exposures on the transcriptome by RNA-seq. Transcriptional profiling was performed on livers from individual male and female mice from the eight exposure groups, using a minimum of N=3 for each sex, with the litter counted as the N. Each group of exposed animals were compared to age-and sex-matched vehicle control animals using a minimum of N=3 per sex/age as detailed in Supplementary Materials and Methods. This profiling revealed exposure-specific, persistent disruption of the transcriptome, and identified vulnerabilities exploited by multiple T2C exposures. These findings are summarized for both sexes and all eight exposures in the bar graphs in Figure 2a, and all exposure DEGs are listed in the combined signature file in Supplemental Table 1. Representative PCA plots and heatmaps for males exposed to PM2.5-JHU and TCDD and females exposed to PM2.5-JHU and TBT are displayed in Figure 2b; comprehensive PCA plots, heatmaps, and volcano plots for all eight exposures and both sexes are provided in Supplemental Figure 2.

**Figure 2.**
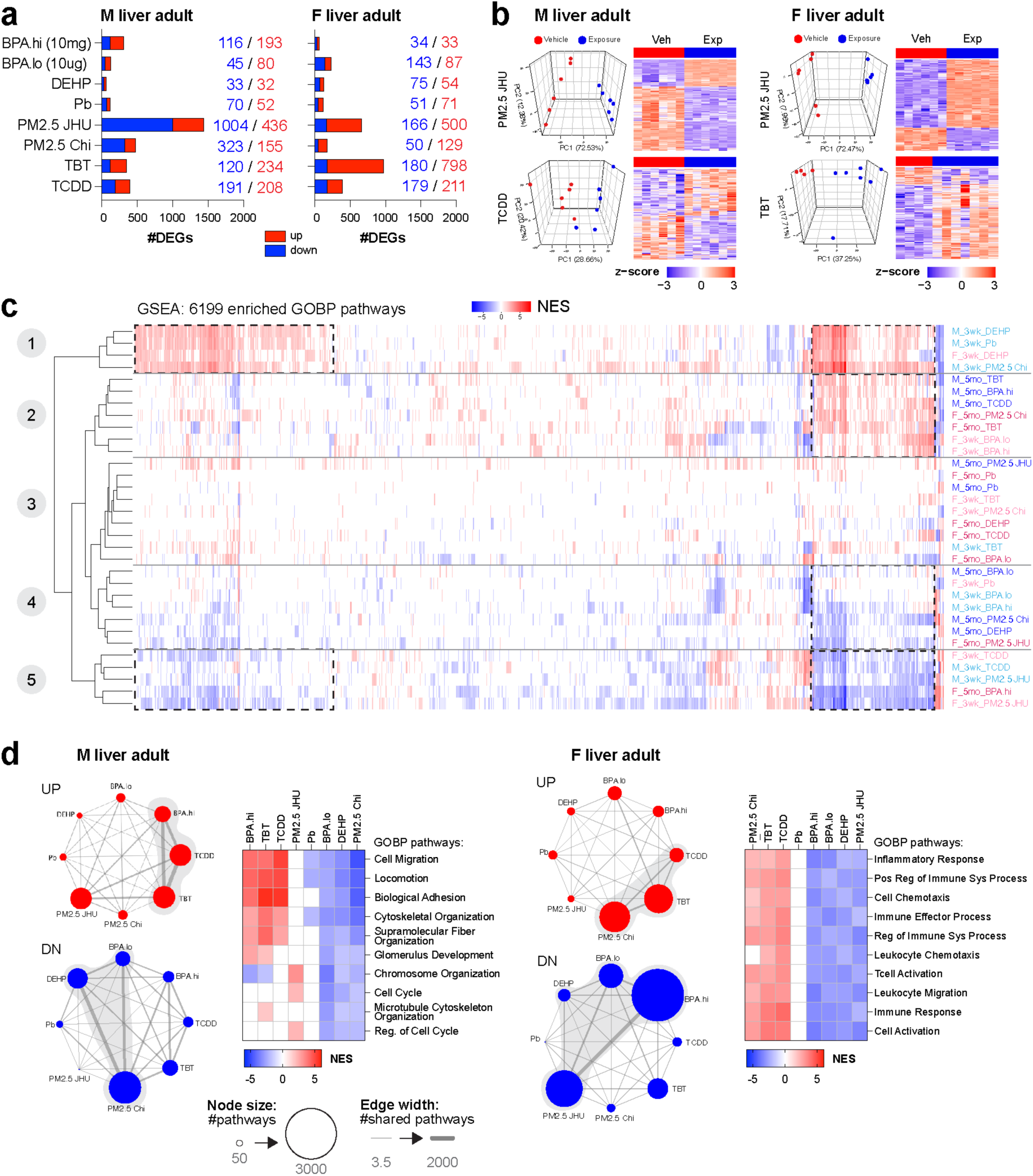
Transcriptome effects of early-life toxicant exposure. a. Summary of the DEGs in male and female mice at 5 months for all 8 exposures. b. PCA and heatmaps for selected female and male mice exposures. c. Enriched pathways via GSEA against GOBP male and female mice, at weanling and at young adult stage, for all 8 exposures. d. Network analysis of overlapping pathways in the same direction across the 8 exposures for male and female mice at 5 months. The node size is proportional to the number of enriched pathways; the edge thickness is proportional to the number of common pathways between two exposures. Overlaps indicated by grey shading are significant (p<10^-100^).

While many early-life exposures induced persistent, long-lasting changes in the adult transcriptome, with numerous DEGs in their exposure signatures, this was not universally the case. The exposures eliciting the greatest to least disruption of the liver transcriptome in young adult male mice, as measured by number of DEGs, were PM2.5-JHU > PM2.5-CHI, TCDD, TBT, BPA.hi > BPA.lo, DEHP, Pb. In females, TBT > PM2.5-JHU, TCDD > BPA.lo, BPA.hi, DEHP, Pb, PM2.5-CHI (Figure 2a). There was also sex bias in how the transcriptome responded to several T2C exposures. For instance, as illustrated in Figure 2a, BPA.hi exposure induced >300 DEGs in livers of male mice but <70 DEGs in livers of exposed females, which were often littermates and therefore equivalently exposed. To assess potential for functional impact of this disruption in the transcriptome, we performed GSEA on the transcriptomes of weanling and young adult mice (Figure 2c and Supplemental Table 2, which provides a complete list of enriched pathways and core genes). We found numerous pathways that were enriched, but remarkably, they clustered into only 5 groups, which was surprising given this extensive data set was derived from mice at different ages (3 weeks and 5 months), both sexes, and in response to eight different exposures conducted at different consortium sites across the country.

Subsequent analysis of the exposures and pathways that defined each of the 5 clusters offered valuable insights into how these early-life environmental exposures had shaped the transcriptome, and revealed common responses to multiple, diverse T2C exposures. Clusters 1 and 5 were primarily defined by pathways enriched in weanlings (Figure 2c). Cluster 1 was characterized by pathways positively enriched in response to DEHP, Pb, and PM2.5, while Cluster 5 was defined by negative enrichment of these same pathways in response to TCDD, PM2.5, and BPA.hi. In contrast, Clusters 2 and 4 were characterized by pathways enriched in young adult mice. Cluster 2 was characterized by pathways positively enriched in response TBT, TCDD, BPA.hi, and PM2.5 and Cluster 4 by negative enrichment of these same pathways in response to BPA.lo, PM2.5, and DEHP. Notably, where the pathways were the same in weanlings and young adults, the exposures were different. As shown in the hatched boxes in Figure 2c, the exposures responsible for positive enrichment of the same pathways in weanlings and young adults in Clusters 1 and 2, respectively, were different, as were the exposures responsible for the negative enrichment of these same pathways in weanlings and young adults in Clusters 5 and 4, respectively. Thus, this surprising convergence revealed using unbiased clustering of pathways from different ages-, sex-, and exposure-specific signatures, suggested that only a limited repertoire of pathways were vulnerable to disruption, becoming either activated or repressed in the liver as a function of exposure, age, and sex of the animals.

To investigate common targets for disruption by these early life exposures, we conducted network analyses to identify those pathways most vulnerable to disruption by T2C exposures—specifically, pathways enriched in response to multiple different exposures. The network diagrams in Figures 2d illustrate the number of enriched pathways (indicated by node size) and the overlaps between pathways enriched in response to the various exposures (represented by thick edges connecting the nodes). In young adult male livers, the nodes corresponding to BPA.hi, TCDD, and TBT (Cluster 2 exposures, Figure 2c, left) shared numerous positively enriched pathways, as indicated in the accompanying normalized enrichment score (NES) heatmap. These pathways included those related to cytoskeletal and fiber organization, adhesion, and locomotion (Figure 2d). Conversely, negatively enriched pathways exhibited strong overlaps between PM2.5-CHI, BPA.Lo, and DEHP (Cluster 4 exposures), of these same pathways (Figure 2d). Similar trends were observed in adult female liver, as depicted in the network diagrams and heatmaps in Figure 2d, right. In female mice exposed to PM2.5-CHI, TBT, and TCDD, young adult livers displayed positive enrichment of pathways associated with immune response, inflammatory signaling, and cell migration, while conversely, livers from female mice exposed to BPA.Hi, BPA.lo, DEHP, and PM2.5-JHU showed negative enrichment of these same pathways.

Thus even for the same exposure, pathway enrichment often exhibited sex specificity. The most significantly enriched pathways in exposed females involved processes for immune cell activation, while pathways related to adhesion and fiber organization were most enriched in the exposure signatures of males. A more detailed TaRGET II examination of how sex influenced responses to all exposures will be presented in a forthcoming Consortium manuscript. Thus, a striking finding from this analysis was that the signatures of diverse T2C exposed livers appeare to be enriched for many of the same pathways, with the direction of enrichment (positive or negative) being exposure-, sex-, and age-specific. To confirm this, we used permutation testing, which confirmed the significance of the shared pathways in the four networks from Figure 2d. This was followed by fitting a normal distribution and determining the z-score for all overlaps-those shown with gray shading in Figure 2d were highly significant (p<10^-100^).

### Multiple Early-Life Exposures Disrupt the Aging Trajectory of Target Genes

The observed overlap between pathways disrupted by T2C exposures shown in Figure 2 was striking given the diverse mechanisms of action of these environmental exposures. This suggests that while the mechanisms of action of T2C environmental toxicants differ significantly, their ability to induce persistent changes in the transcriptome converged on a discrete, shared, set of vulnerabilities. To identify these shared vulnerabilities, we compared the 16 T2C exposure signatures from adult male and female livers with composite aging signatures derived from normal 3-week and 5-month T2C male and female livers (https://dcc.targetepigenomics.org/). This analysis revealed that liver agingDEGs constituted a remarkable fraction of the T2C exposure signatures, accounting for approximately 40-60% DEGs in exposed male and 30-60% in exposed female signatures (Figure 3a and Suppl Figure 3a). In males, the agingDEG component of the exposure signatures ranged from a low of 38% (PM2.5-JHU) to a high of 60-62% (BPA.hi and DEHP) and 43-53% of the signatures for TBT, TCDD, BPA.lo, PM2.5-Chi, and Pb. While agingDEGs made up a significant proportion of all exposure signatures, only a fraction of the 2,582 agingDEGs identified in males, for example, were disrupted by one or more of the T2C exposures. This indicates that the observed exposure-induced changes in agingDEG expression were not the result of a global change in how the liver itself had aged, as fewer than 20% of all aging DEGs were reprogrammed in response to T2C exposures. Additionally, within exposure groups, we observed both increases and decreases of aging DEGs, i.e. acceleration and attenuation of the normal trajectory of age-associated gene expression, also indicating no organ-level shift in biological aging. Rather, this reprogramming of agingDEGs by multiple exposures suggests that these genes were a vulnerable target, perhaps because they retained the plasticity to change expression as a function of age, rendereing them susceptible to reprogramming by environmental exposures.

**Figure 3.**
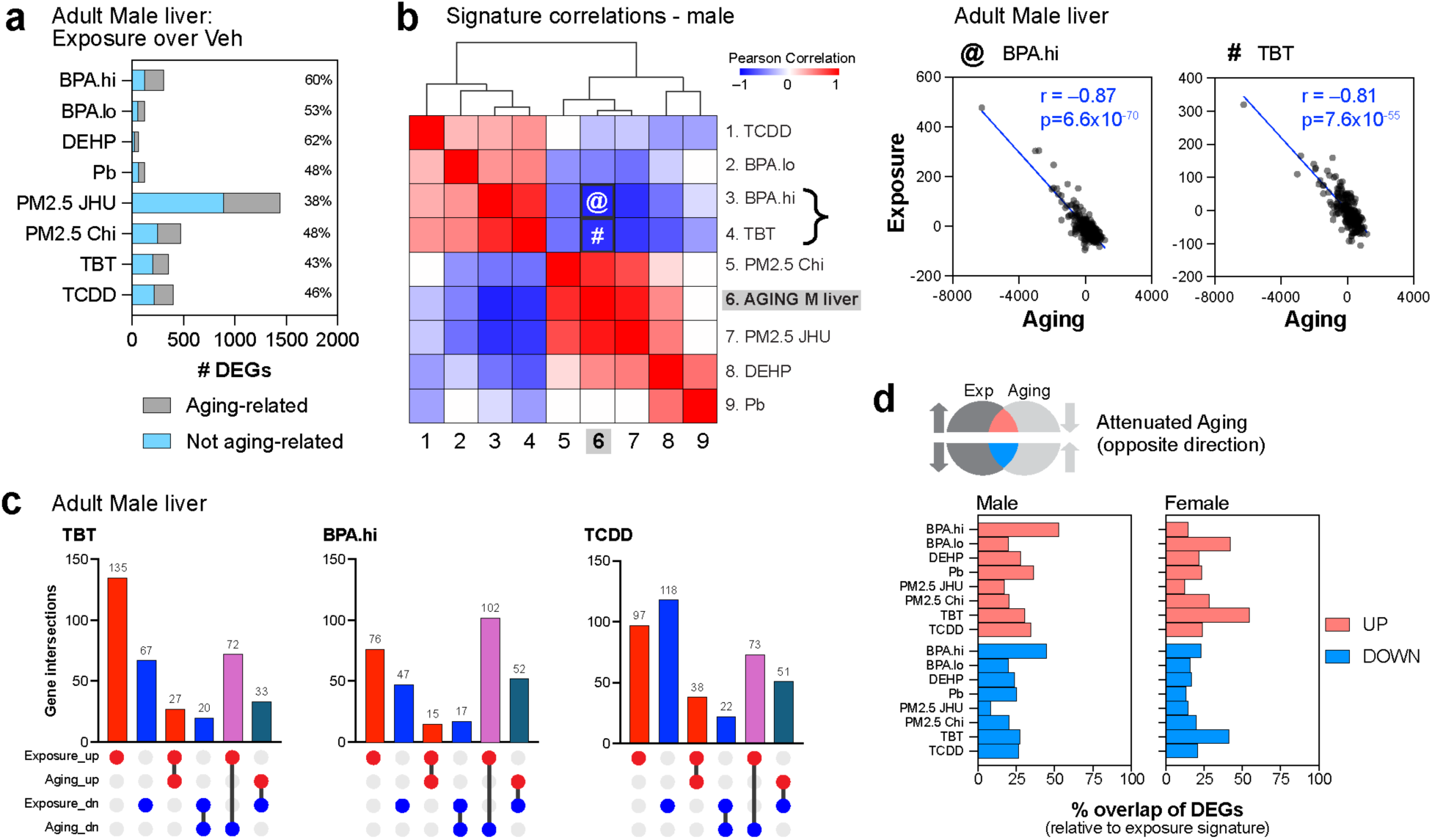
Robust overlap between exposure response and aging DEGs is seen in both in male and female mice. a. Bar graph showing the overlap between exposure DEGs and aging DEGs in males irrespective of direction of change. b. Matrix of Pearson’s correlation coefficients (p<0.05) between summed z-scores for exposure exposure signatures and agingDEGS in male mice over the GTEx liver transcriptome dataset. c. UpSet plots showing overlap between aging DEGs and selected exposures TBT, BPA, TCDD in male mice. d. Summary bar chart showing the proportion of exposure DEGs associated with attenuated aging in males and female

An overview of the reprogramming of the normal trajectory of agingDEGs is provided in the Pearson correlation matrices against the GTEx normal liver transcriptome, using male (Figure 3b) and female (Supplemental Figure 3b) datasets. This analysis highlights those T2C exposure signatures that were strongly and inversely correlated with normal aging, indicating an attenuation of typical gene expression changes that would normally occur between weaning and young adulthood. For example, in the adult male liver, the BPA.hi exposure signature exhibited a strong negative correlation (r =-0.87) with normal aging, indicating reprogramming attenuated the trajectory of target agingDEGs, as did TBT (r =-0.81) (Figure 3b). In females (Supplemental Figure 3b), TBT exposure resulted in an even stronger negative correlation with normal aging (r =-0.99), as did PM2.5-JHU (r =-0.79), indicating both these exposures strongly attenuated the aging trajectory of signature agingDEGs. UpSet plots showing the change in trajectory of agingDEGs in the transcriptional signatures for TBT, BPA.hi, and TCDD in males are shown in Figure 3c, and a list of the agingDEGs in their exposure signatures showing the greatest change in expression are provided in Table 2; the complete list of agingDEGs for all exposure signatures can be derived from the combined signatures in Supplemental Table 1. UpSet plots provided in Supplemental Figure 3c and 3d highlight how the normal trajectory of agingDEGs shifted (accelerated or attenuated) in response to early-life exposures.

**Table 2.**
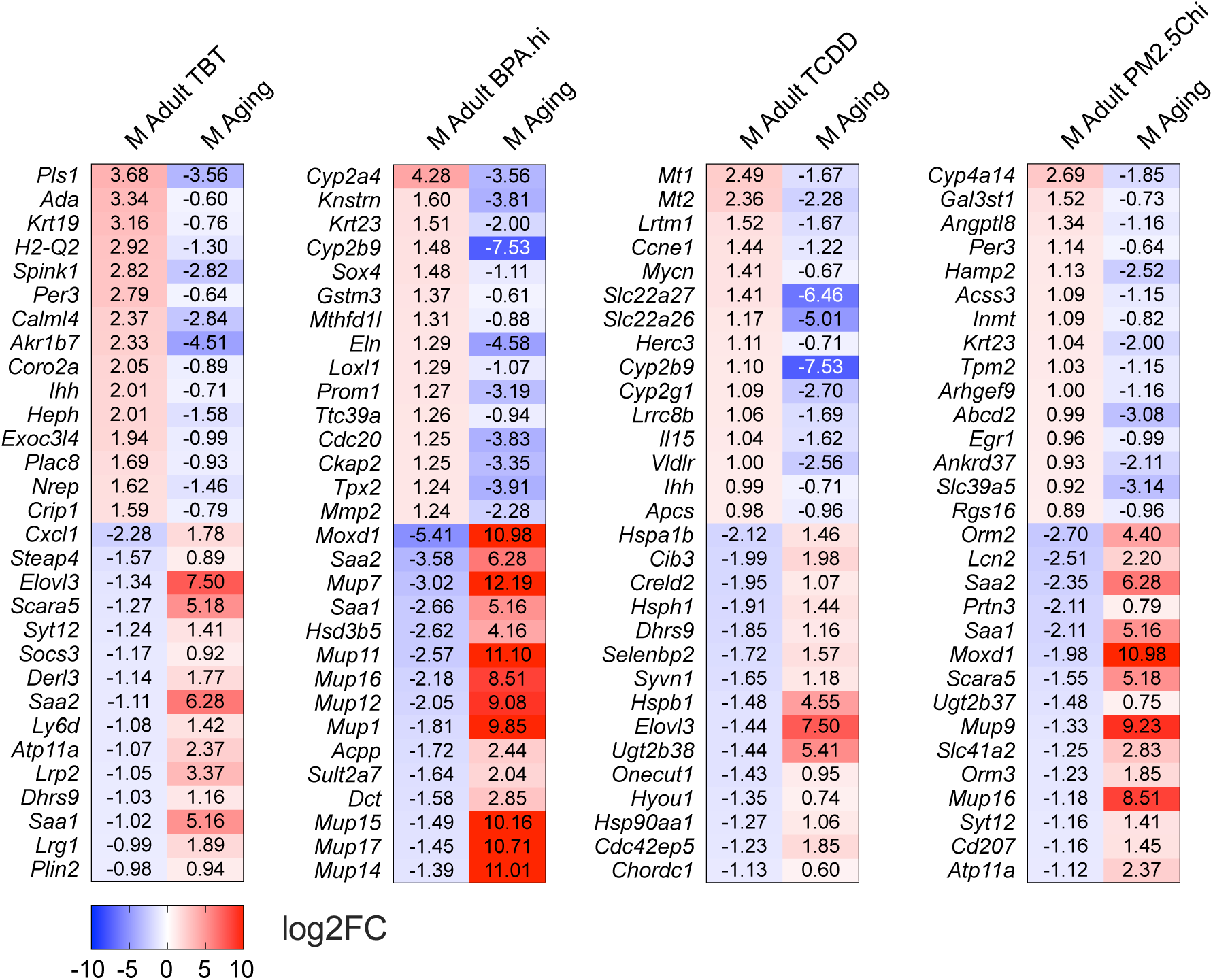
Top aging DEGs in all exposure signatures.

From these analyses, a pattern of attenuated agingDEG expression emerged as a shared response to several T2C exposure, where agingDEGs that normally increased with age failed to do so (i.e. remained low) and agingDEGs that typically decreased, remained high. Venn diagrams in Supplemental Figure 3e depict the overlap between signature DEGs and agingDEGs across all exposures highlighting this feature of attenuated aging. Figure 3d provides a summary of how early-life T2C exposures attenuated the aging trajectory of target agingDEGs in the adult liver of both males and females. These findings suggest that, despite distinct modes of action, reprogramming of the transcriptome by T2C exposures converged on a shared biological vulnerability: the inherent plasticity of target genes to change expression with age.

### Histone Modifications Reveal Epigenomic Aging Patterns in the Male Mouse Liver

Histone modifications, acting as transcriptional ‘rheostats’ to finely tune gene expression, have emerged as key targets of environmental exposures ^2,19^. However, the dynamics of histone modifications as a function of age remain less understood compared to the plasticity of other epigenetic mechanisms, such as DNA methylation ^19–21^. Thus, to characterize the impact of early-life exposures on the epigenome during the T2C study interval, we first performed ChIP-seq profiling for active (H3K27ac, H3K4me1, H3K4me3) and repressive (H3K27me3, H3K9me3) histone marks during normal aging across the T2C study interval. These data were used to map chromatin states genome-wide, including promoters and enhancers (H3K4me3, H3K27ac, H3K4me1), as well as facultative (H3K27me3) and constitutively repressed (H3K9me3) chromatin. These TaRGET II consortium-wide epigenomic profiling studies were limited to males, selected because males displayed the strongest phenotypic response to several T2C exposures, such as the liver tumors observed in males in response to TBT ^22,23^. During the time from weaning until 10 months of age, >350 epigenomic profiles for histone modifications were generated from individual vehicle control and exposed male mice.

Differential enrichment of these histone modifications was determined in the male liver over two time-intervals: from weaning to early adulthood (3 weeks to 5 months) and from early to later adulthood (5 to 10 months). This profiling identified differentially enriched peaks (DEPs) in the liver as animals aged (agingDEPs) from weaning to early and later adulthood. For active histone marks H3K27ac, H3K4me1, and H3K4me3, robust remodeling of the epigenome occurred during the transition from weaning to early adulthood, as illustrated by the Circos plots and bar graphs in Figure 4a, and the heatmaps for individual marks in Supplemental Figure 4a. During this period, both increases and decreases in these marks occurred, although loss of active marks predominanted as weanlings aged into young adults. Notably, reduction in active histone marks aligns with the widespread decrease in gene expression observed in the liver transcriptome during this time (Figure 1b). In the subsequent transition from early to later adulthood, the loss of H3K27ac and H3K4me3 marks predominated, while H3K4me1 remained relatively unchanged (Figure 4a, Supplemental Figure 4a).

**Figure 4.**
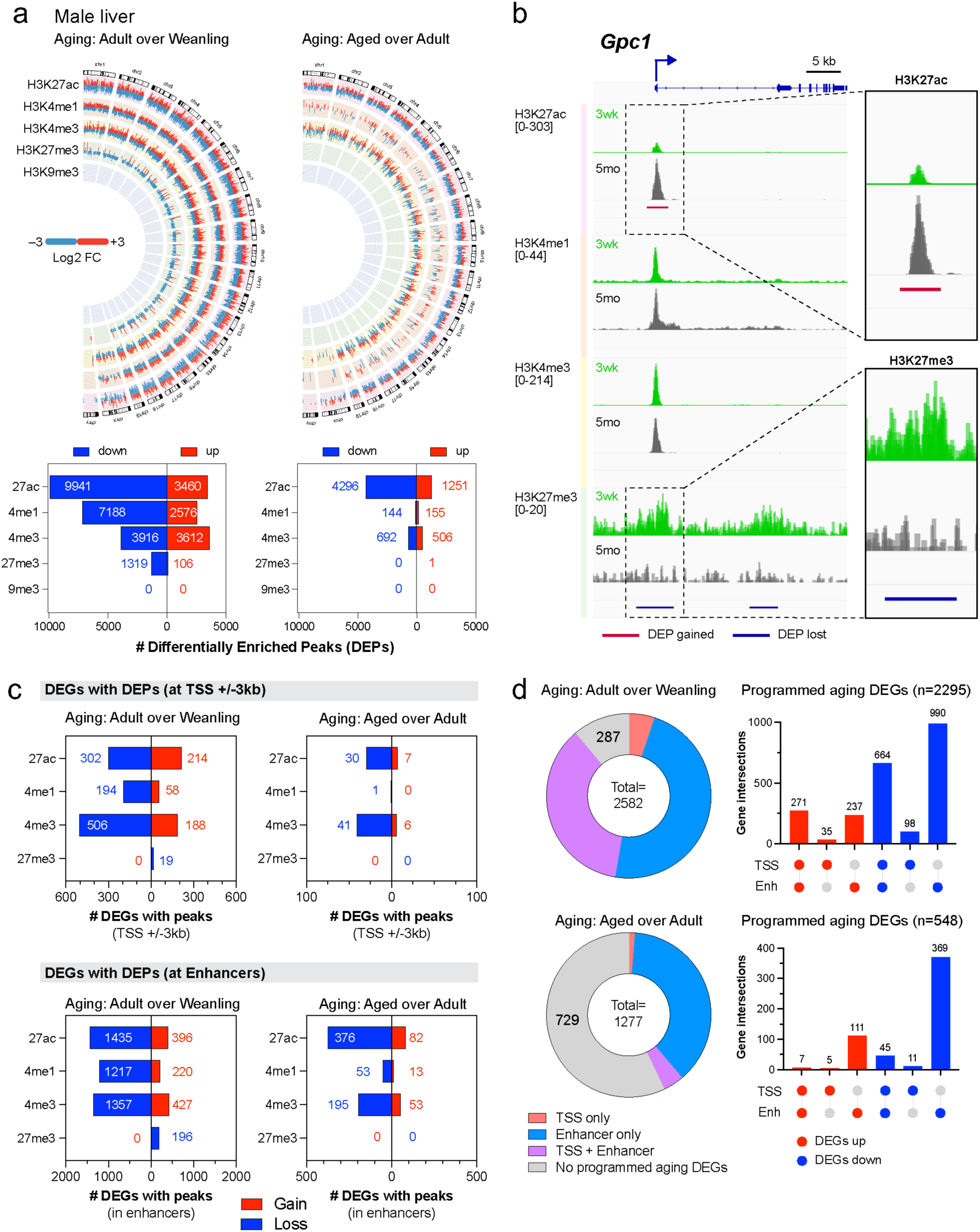
Epigenome programming associated with aging from weanling to young adult to aged adult. a. Circos plots indicating direction and magnitude of changes in histone marks with age across the genome, and summary bar graphs. b. IGV browser indicating aging DEPs and histone marks signal at the Gpc1 locus. c. Bar graphs showing a summary of aging DEGs concordant with agingDEPs at promoters (TSS +/-3Kbp) and enhancers. d. UpSet plots and pie charts of combinatorial concordant agingDEP programming at TSS and enhancers for agingDEGs.

The dynamics of repressive histone marks showed patterns distinct from active marks. H3K9me3, a cell-specific constitutive repressive mark, exhibited minimal change across either age interval (Figure 4a), consistent with the important role of this mark in silencing lineage-inappropriate gene expression and maintaining cell identity in the mature liver. H3K27me3, a facultative repressive mark, demonstrated broad clustering across differentially enriched regions, which contrasted to the sharp peaks characteristic of other marks (Figure 4b). H3K27me3 exhibited a sharp decline between weaning and early adulthood but was changed little between early and later adulthood (Figure 4a). This age-related decline in H3K27me3 is consistent with previous findings ^24^. However, the reported gain in that study of large H3K27me3-enriched domains in much older mice (70-95 weeks), was not observed within the 10-month timeframe of the T2C longitudinal study, as illustrated in the IGV tracks in Figure 4b as consistently observed in vehicle control animals from two T2C sites where ChIP-seq was conducted through 10 months (other sites contributed livers only to the 5-month timepoint). These findings suggest that in the male liver, early loss of H3K27me3 precedes the later emergence of broad enriched domains, and the shift to what has been termed a’hyper-repressive state’ ^24^ occurs later than the 10-month window for the T2C study.

We also profiled genome-wide changes in histone marks at promoter and enhancer regions as a function of age, as detailed in Supplemental Figure 4b and Supplemental Table 3. To ensure precise linkage between genes and the enhancers known to regulate their expression, we utilized the enhancer-gene pairs cataloged by Enhancer Atlas ^25^. To do so, we first defined enhancer’anchors’ (Supplemental Figure 4c), where enhancers identified across different studies and tissues, but located adjacently, were consolidated into a single’enhancer anchor’ (Supplemental Table 4). Genes potentially regulated by these enhancer anchors were then identified using the Enhancer Atlas compendium.

While there were some gains in histone marks between weaning and early adulthood, the predominant change was a loss of active marks in both promoter – defined as transcription start site (TSS) of genes ±3kb – and enhancer regions. Notably, however, compared to promoters, nearly an order of magnitude more agingDEPs mapped to enhancers. Between 5 and 10 months, further loss of active marks was observed at both promoters and enhancers, with enhancer regions again exhibiting significantly more reprogramming compared to promoters (Supplemental Figure 4b and Supplemental Table 3).

Enhancers are an order of magnitude more abundant than genes (ENCODE 2012), implying a “many-to-one” interaction structure where many different enhancers regulate a single gene. Previous studies have shown less than only 1% of enhancers are highly pleiotrophic (i.e. regulate multiple genes), 50% of enhancers are linked to only a single gene, and the average number of genes linked to a single enhancer is ∼2.5 ^26^. Our analysis of enhancer anchors aligned well with this prediction: we found on average enhancer anchors with agingDEPs were linked to 1.8 agingDEGs. In contrast, while each enhancer was linked to only a few genes, the number of enhancers that can regulate any single gene may be quite large. This is thought to help stabilize or “buffer” gene expression, and provide robust enhancer plieotrophy across multiple tissues ^26^. In our stady, on average, agingDEGs were linked to ∼8 linked enhancer anchors with agingDEPs. Thus, the predominance of agingDEPs occurring at enhancers vs promeotrs may be due to the fact that the merging enhancers identified in multiple tissues into a single enhancer anchor captures this plieotrophy, while any single agingDEG was asigned only a single promoter region (TSS +/-3 kb).

We next assessed the relationship between the aging epigenome and aging transcriptome by mapping agingDEPs for H3K4me3, H3K27ac, H3K4me1, and H3K27me3 to promoter regions and linked enhancers of agingDEGs-H3K9me3, which was unchanged s a function of age, was not included (Supplemental Table 3). Consistent with expectations, during the transition from weaning to young adulthood, agingDEGs whose expression decreased showed loss of active histone marks (H3K27ac, H3K4me1, and H3K4me3) or gain of H3K27me3 at promoters and enhancers, while agingDEGs whose expression increased showed gains of active marks at associated regulatory regions, or in a few cases, loss of H3K27me repressive marks (Figure 4c). During this interval, agingDEPs at enhancers linked to agingDEGs were considerably more frequent compared to agingDEPs at promoter regions of agingDEGs (Figure 4c). In contrast, during the transition from early to later adulthood, fewer agingDEGs were associated with agingDEPs. Although we identified 619 downregulated and 658 upregulated agingDEGs between 5 and 10 months of age (Supplemental Table 3), fewer than 100 DEPs occurred at the promoter regions of these agingDEGs, and fewer than 400 at their linked enhancers (Figure 4c), i.e less than half were associated with agingDEPs at promoters/enhancers.

To understand combinatorial changes in epigenetic histone marks that occurred with aging, we also generated UpSet plots for H3K27ac, H3K4me1 and H3K4me3 DEPs at the promoters and linked enhancers of agingDEGs (Supplemental Figure 4d) during the transition from weaning to young adulthood and young to later adulthood. This analysis revealed that H3K27ac and H3K4me3 were the primary histone marks associated with programmed epigenomic aging. Between weaning and early adulthood, loss of H3K4me3 alone was the predominant change at promoters of agingDEGs whose expression decreased, and although agingDEGs whose expression increased during this interval were fewer, these agingDEGs gained H3K4me3 and H3K27ac, alone or in combination, at their promoters. Notably, H3K4me1 showed little change at promoters of agingDEGs.

At linked enhancers, coordinated loss of all three H3K27ac, H3K4me1 and H3K4me3 marks was the primary change for agingDEGs whose expression decreased. Enhancers linked to agingDEGs whose expression increased showed coordinate gain of all three H3K27ac, H3K4me,1 and H3K4me3 marks, or H3K27ac and H3K4me3 (without H3K4me1). Between early and later adulthood, fewer DEPs were mapped to promoters and linked enhancers, and during this time, gains/losses of H3K27ac, with or without changes in H3K4me3, was the predominant change observed.

To obtain a more precise estimate of the overall association between concordant changes in the aging transcriptome and aging epigenome, we employed UpSet plots to classify agingDEGs and agingDEPs as 1) near the promoter (TSS ± 3 kb), 2) at a linked enhancer, 3) in both regions, or 4) in neither where decreased expression correlated with the loss of active or gain of repressive marks and increased expression correlated with gain of active or loss of repressive marks. As illustrated in the UpSet plots and pie charts in Figure 4d, all but 287 of the 2,582 male agingDEGs identified between weaning and early adulthood exhibited concordant changes in epigenetic histone marks in at least at one associated regulatory element (over 88% concordance). During the transition from early to later adulthood, a period with fewer agingDEGs, only 42% of the 1,277 agingDEGs were associated with agingDEPs in either their promoter regions or linked enhancers. These findings indicate extensive concordance between the male liver aging transcriptome and the programmed aging of epigenetic histone marks, especially during the interval from weaning to young adulthood.

Finally, to identify functional and cell type-specific correlates between the aging epigenome and transcriptome, we performed Over-Representation Analysis (ORA) using the specific subset of agingDEGs with concordant agingDEPs in their promoter region or linked enhancers, as shown in Supplemental Figure 4e. AgingDEGs that lost active histone marks and exhibited decreased expression were enriched for nonparenchymal cell identity genes, whereas agingDEGs whose expression increased and gained active marks were enriched for hepatocyte cell identity genes (Supplemental Figure 4e). This aligns with the cell-type-specific transcriptional shifts in Figure 1e, where hepatocyte identity genes increased in expression with age, while nonparenchymal cell identity genes showed a decrease. Additionally, ORA using the Reactome compendium revealed that pathways positively enriched with upregulated agingDEG/DEPs were involved in metabolism, while pathways negatively enriched with downregulated agingDEG/DEPs were associated with cellular architecture and extracellular matrix production (Supplemental Figure 4e and Supplemental Table 5). These findings indicate that, at both gene and pathway levels, age-related epigenomic programming of histone marks is highly concordant, and aligns well with the direction and cell-type specificity of changes observed in the aging transcriptome of the male mouse liver.

### Early-Life Exposures Reprogram the Adult Epigenome

Mapping of exposure-induced DEPs (Figure 5a) to gene promoter (Supplemental Table 6) and enhancer (Supplemental Table 7) regions allowed us to link changes in the transcriptome to epigenomic reprogramming induced by early-life T2C exposures in male mouse livers. Interestingly, the epigenomic states (promoters vs enhancers), extent of the reprogramming (number of DEPs), as well as the specific histone marks affected, varied dramatically across the different exposures. As a result, T2C exposures that elicited persistent reprogramming were each associated with unique epigenomic signatures. as summarized in the bar charts for all eight exposures in Figure 5a and visualized in heatmaps in Supplemental Figure 5a. For example, PM2.5-Chi predominantly induced gains in H3K4me3, with minimal changes in H3K27ac or H3K4me1. In contrast, TBT exposure caused both gains and losses of H3K27ac and H3K4me3, with losses prevailing, while leaving H3K4me1 largely unaffected. TCDD exposure caused a net loss across all three active marks – H3K27ac, H3K4me3, and H3K4me1. In the case of BPA.hi, DEPs were primarily gains in H3K27ac and H3K4me1, with minimal impact on H3K4me3. Notably, this variability in how T2C exposures reprogrammed the epigenome contrasted with their effect on the transcriptome, where convergence on a discrete set of pathways was seen (Figure 2), further reinforcing that despite their different mechanisms of action, and different exposures signatures, their convergent impact on the transcriptome was due to targeting common vulnerabilities.

**Figure 5.**
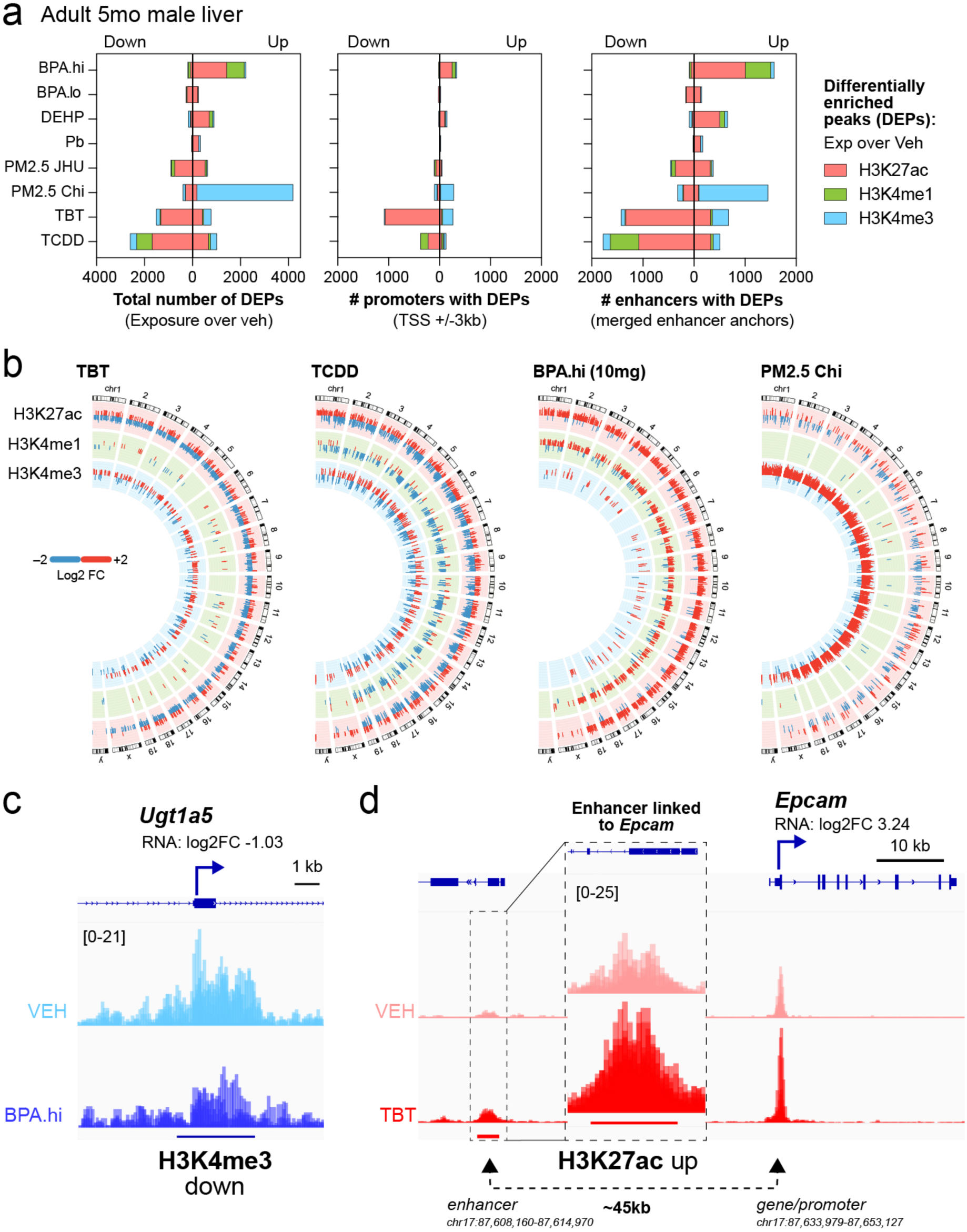
Epigenome reprogramming associated with early-life toxicant exposures. a. Summary bar graphs for exposure-induced DEPs and their overlap with promoters and enhancers. b. Circos plots showing direction and magnitude for epigenomic reprogramming by TBT, TCDD, BPA.hi, and PM2.5-Chi. c. Selected IGV tracks.

As seen with aging, persistent epigenomic reprogramming induced by these exposures was more robust at enhancers than at promoter regions for their target genes (Figure 5a). For instance, as shown in Figure 5a, early-life TCDD exposure led to loss of histone marks at ∼300 promoter regions but >1800 enhancers in early adulthood. This disparity is unlikely to be stochastic, as the cumulative target sizes for promoter regions (±3kb from the TSS) vs enhancer anchors (where any DEP was required to map within the enhancer anchor itself) were 131 Mb and 326 Mb, respectively. Possible explanations for the observed preference for enhancer vs promoter reprogramming is that exposures preferentially target enhancers compared to promoter regions (i.e, enhancers are more susceptible to reprogramming) or that when reprogrammed, DEPs in enhancers are more enduring than those in promoter regions (i.e. enhancer reprogramming is more persistent).

These findings underscore that in the male mouse liver, early-life T2C exposures drive long-lasting, exposure-specific epigenomic reprogramming in adulthood, varying markedly in both the number, direction, and specific epigenetic marks affected. The variability in epigenomic reprogramming is further illustrated with Circos plots for TCDD, TBT, BPA.hi, and PM2.5-Chi (Figure 5b), and other exposures in Supplemental Figure 5b. However, while some early-life T2C exposures induced robust changes in the adult liver epigenome, others had minimal effect. T2C exposures ranked for extent of reprogramming (as determined by number of DEPs), were PM2.5-Chi > TCDD, TBT, and BPA.hi > PM2.5-JHU and DEHP > Pb and BPA.lo (Supplemental Tables 6 and 7).

Importantly, the persistent epigenomic reprogramming induced by T2C exposures was not merely a consequence of altered gene expression in the exposed male livers, as extensive reprogramming also occurred in promoter regions and linked enhancers of genes where expression was unchanged, a phenomenon previously referred to as “silent reprogramming” ^27^. This is evident from the disproportionate number of reprogrammed promoters and enhancers relative to exposure-induced DEGs for several exposures. For example, TBT exposure resulted in the loss of H3K27ac at over 1,000 promoters and more than 1,200 enhancers (>2,200 DEPs), yet its adult liver exposure signature comprised only 354 DEGs, with just 120 being downregulated (compare Figure 5a and Figure 2a). Similarly, BPA.hi increased H3K27ac and H3K4me1 at more than 200 promoters and over 1,500 enhancers (>1,700 DEPs), but was associated with only ∼300 DEGs. PM2.5-Chi exposure induced gains in H3K4me3 at over 200 promoters and more than 1,500 enhancers (>1,700 DEPs) but resulted in an exposure signature of just 478 DEGs. This “silent reprogramming” mirrors findings from perinatal BPA exposure in rats ^27^, where early-life BPA “primed” genes for aberrant transcriptional responses to later-life stressors. In that study, epigenetic reprogramming of EGR1 target genes by BPA led to exaggerated transcriptional activation under high-fat diet conditions, ultimately causing metabolic dysfunction. These results suggest that even the silent epigenomic changes observed in response to T2C exposures could serve as a latent mechanism for maladaptive responses to other stressors later in life.

To link signature DEGs to exposure-induced epigenomic reprogramming, we mapped exposure-induced DEPs to the promoter and enhancers regions for each DEG in the male exposure signature (Supplemental Table 8). UpSet plots summarizing the overlap between DEGs and DEPs in promoter or enhancer regions are shown in Supplemental Figure 5c and d. Between 25-33% of promoters and/or linked enhancers of signature DEGs had undergone epigenomic reprogramming in livers exposed to BPA.hi, TBT, TCDD, and PM2.5-Chi (Table 3). This association became even more pronounced when the direction of the global epigenomic response (Figure 5a) was aligned with the direction of DEG expression. For instance, BPA.hi primarily induced genome-wide gain in active marks such as H3K27ac, H3K4me3, and H3K4me1, and 53% of BPA.hi DEGs that increased were linked to gains in these marks at their promoters and/or enhancers. Similarly, TBT predominantly drove loss of H3K27ac and H3K4me1, with 51% of DEGs exhibiting decreased expression associated with corresponding losses in these marks. In contrast, exposures with minimal impact on the epigenome, PM2.5-JHU, BPA.lo, DEHP, and Pb, exhibited the fewest signature DEGs and only limited epigenomic reprogramming in promoter or linked enhancer regions (5–16%; Table 3).

**Table 3.** Promoter and/or Enhancer Reprogramming at T2C Signature DEGs.

| Exposure | Signature DEGs | DEGs with DEPs | Percent | Aging DEGs overlap | Aging DEPs overlap | Percent |
| --- | --- | --- | --- | --- | --- | --- |
| TBT | 354 | 122 | 34% | 152 | 50 | 33% |
| TCDD | 399 | 122 | 31% | 184 | 50 | 27% |
| BPA.hi | 309 | 114 | 37% | 186 | 74 | 40% |
| PM2.5Chi | 478 | 122 | 26% | 229 | 57 | 25% |
| PM2.5Bal | 1440 | 237 | 16% | 545 | 87 | 16% |
| BPA.lo | 125 | 18 | 14% | 66 | 8 | 12% |
| DEHP | 62 | 9 | 15% | 40 | 4 | 10% |
| Pb | 122 | 6 | 5% | 37 | 1 | 3% |

This relationship extended to agingDEGs reprogrammed by BPA.hi, TBT, TCDD, and PM2.5-Chi. As shown in Table 3, 25–40% of agingDEGs in these exposure signatures displayed epigenomic reprogramming at promoters and/or linked enhancers, further underscoring the interplay between exposure-induced epigenetic alterations and persistent reprogramming of the adult transcriptome (Supplemental Table 8). Examples of this reprogramming are illustrated in the IGV tracks for *Ugt1a5*, which lost H3K4me3 in the promoter and was decreased in the BPA.hi signature (Figure 5c), and *Epcam*, which gained H3K27ac in its linked enhancer, and was increased in the TBT signature (Figure 5d). Additional examples are provided in Supplemental Figure 5e-f for genes reprogrammed and differentially expressed in response to PM2.5.Chi, TCDD, and TBT. As shown in Supplemental Table 8, whereas agingDEPs were identified in ∼8 linked enhancers/agingDEG, fewer linked enhancers were reprogrammed per signature DEG. Interestingly, for BPA.hi, PM2.5.Chi, TBT, and TCDD, where agingDEGs made up the largest portion of their exposure signatures, the same enhancers associated with programmed aging were also the target for reprogramming for 40-67% of these agingDEGs, further highlighting that the plasticity inherent in epigenetic aging represents a vulnerability exploited by multiple exposures. Taken together, the picture that emerged for BPA.hi, TBT, TCDD, and, to a lesser extent, PM2.5-Chi, was that despite differences in their modes of action, these early-life exposures induced significant epigenomic reprogramming, most prominently at enhancers linked to their signature DEGs, further implicating enhancer-driven mechanisms as a key feature of the long-lasting effects of early-life exposures on programmed aging. Thus enhancer reprogramming may play a larger, and previously unappreciated, role than promoters in reprogramming by early life exposures.

### Early-Life Exposures Polarize the Adult Transcriptome with Cell-type and Directional Specificity

In the liver, metabolically active hepatocytes constitute approximately two-thirds of cells, while non-parenchymal cells, such as Kupffer, Stellate, and sinusoidal endothelial cells, make up the remaining third, each contributing to liver structure and function in unique ways ^28^. To explore how exposure-induced reprogramming impacts specific liver cell types, we conducted GSEA on the transcriptomes of both male and female mice using identity genes for epithelial parenchymal cells (hepatocytes and bile duct epithelial cells/cholangiocytes) and various non-parenchymal cell types, including Stellate, Kupffer, endothelial, and resident immune cells (B-cells and NK-NKT cells). This analysis revealed a remarkable polarization in the exposure signatures of TBT, TCDD, BPA.hi, and PM2.5-Chi, with pathway enrichment for liver cell identity genes (positive or negative) for all four exposures exhibiting striking directionality and distinct relationship to normal aging.

Identity genes for the 32 liver-resident cell types ^29^ (Supplemental Figure 6a) were grouped into two parenchymal (hepatocytes and bile duct epithelial) and five non-parenchymal (Stellate, Kupffer, NK, vascular, and B cells) categories, based on their collective identity genes (Figure 6a, males and Supplemental Figure 6b, females). Although we anticipated cell-type specificity in gene expression changes caused by T2C exposures, an unexpected finding was the clear directionality, and pronounced anti-correlation with programmed aging, observed across all cell types in both male and female livers. In males exposed to TCDD, TBT, BPA.hi, and PM2.5-Chi, hepatocyte agingDEGs that normally increased from weaning to young adulthood exhibited attenuated aging, remaining low relative to age-matched 5-month-old controls—a similar effect was observed in bile duct epithelial cells exposed to PM2.5-Chi (Figure 6a). Remarkably, non-parenchymal cells also showed attenuated aging, but in the opposite direction: agingDEGs that typically decreased from weaning to young adulthood remained high in sinusoidal endothelial, Kupffer, Stellate, and resident immune cells in the male liver relative to age-matched 5-month-old controls in response to TCDD, TBT, and BPA.hi (Figure 6a).

**Figure 6.**
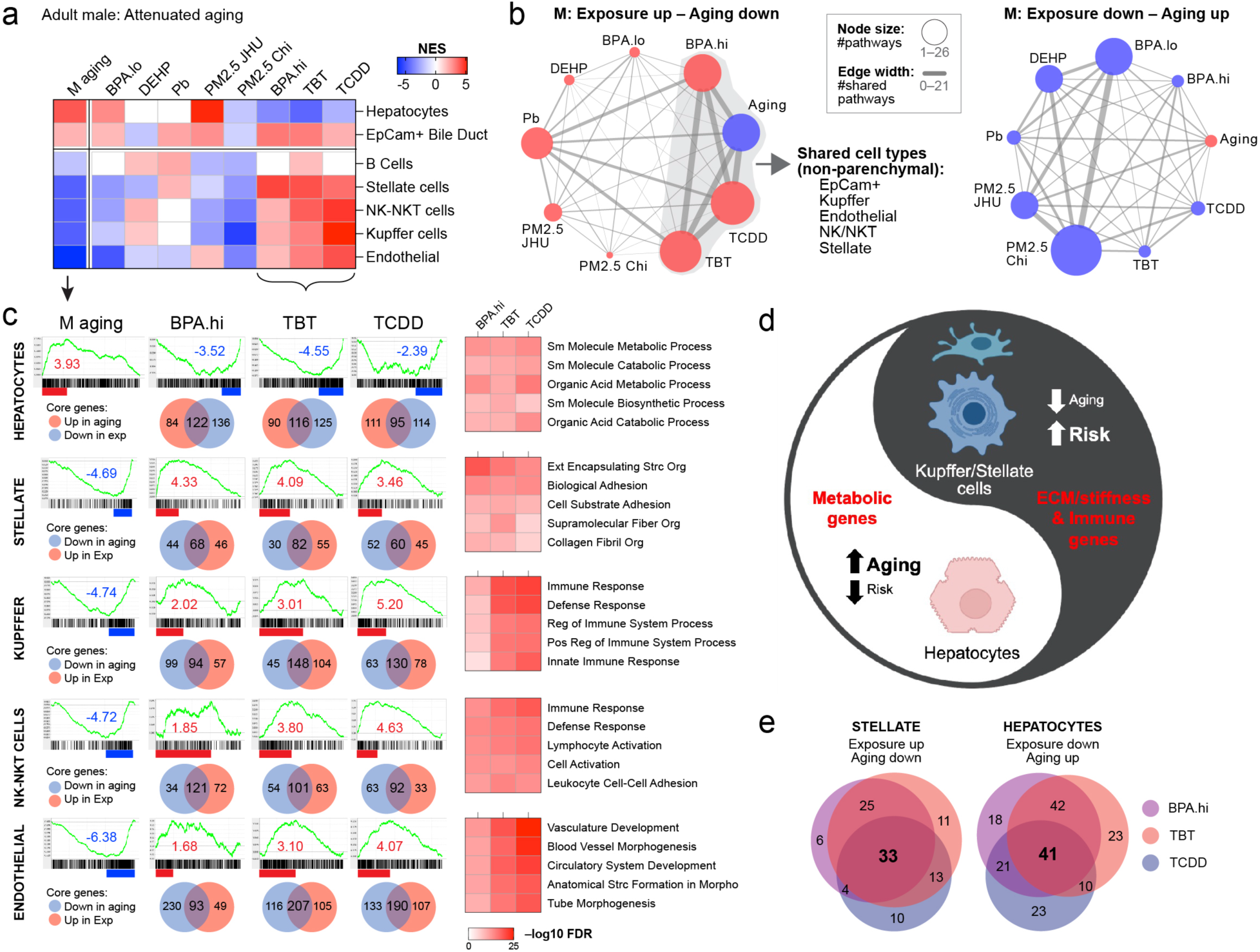
Environmental exposure induced cell type specific attenuated aging polarized effects. a. GSEA enrichment results showing attenuated aging after exposure in 5-month males using 7 summary cell-type signatures from the Aizarani liver compendium. b. Network analysis of cell identity genes vs aging in male livers. c. GSEA enrichment plots with Venn diagrams for overlaps between aging and exposure core genes from BPA.hi, TBT, and TCDD in 5-month males for Stellate, Kupffer, MVECs, LSECs, and Hepatocytes. GSEA core genes were further analyzed using ORA for enriched gene sets from the Reactome and GOBP compendia. d. Inverse relationship between aging processes and cell-type specificity derived from T2C exposure signatures. e. Venn diagrams for overlap of TBT, TCDD, and BPA.hi core genes reprogrammed in hepatocytes and Stellate cells.

In females, a similar attenuated aging effect, characterized by suppressed expression of hepatocyte agingDEGs, was observed in response to PM2.5-JHU, BPA.lo, Pb, TBT, and TCDD. In bile duct epithelial cells, this effect was also evident in response to BPA.hi, PM2.5-JHU, and TCDD. Conversely, as seen in males, non-parenchymal cell agingDEGs that usually declined with age remained high in livers exposed to BPA.lo, Pb, TBT, TCDD, and PM2.5-Chi (Supplemental Figure 6b).

Network diagrams show the extent to which the various non-parenchymal liver cell types exhibited this attenuated aging effect. These diagrams depict the extent of overlap between nonparenchymal cell identity genes that normally change with age and those in the exposure signatures, with edge thickness denoting the degree of overlap between cell identity agingDEGs and exposure DEGs. As shown for males (Figure 6b) and females (Supplemental Figure 6c), there was extensive overlap— albeit in the opposite direction—between agingDEGs and signatures DEGs in multiple non-parenchymal cell types. Notably, however, as described above, this was the case only for those aging DEGs that normally decreased between weaning and adulthood. For example, in males, agingDEGs that typically decrease with age (shown as the blue node in “Exposure up” network diagram in Figure 6b) remain elevated (red nodes) in many non-parenchymal cell types following exposure to BPA.hi, TCDD, and TBT. While there were fewer non-parenchymal cell identity genes that increased with age (note the small red aging node in the “Exposure down” network diagram in Figure 6b), these genes showed little overlap with those in the T2C exposure signatures despite the fact that several signatures contained large numbers of identity DEGs genes that were decreased (note the large blue exposure nodes in that diagram). A similar pattern emerged in females, where the most pronounced reprogramming of non-parenchymal cell types occurred in response to TCDD, PM2.5-Chi, and TBT, resulting in higher expression of agingDEGs that would have normally decreased with age (Supplemental Figure 6c). Thus, in both sexes, reprogramming induced by multiple exposures exhibited dramatic cell-type specificity and directionality, resulting in a polarized attenuated aging signature of increased expression of agingDEGs in nonparenchymal cells and decreased expression of agingDEGs in parenchymal cells.

For the exposures that had the greatest impact, we also conducted GSEA to identify functional pathways normally up-or down-regulated as a function of age in parenchymal and non-parenchymal cells. In males, Figure 6c shows enrichment plots for cell-identity genes that change with age (far left panels) for hepatocytes, stellate, Kupffer, NK-NKT, and endothelial cells, and enrichment plots for the exposure signatures of BPA.hi, TBT, and TCDD (enrichment plots to the right). Taking the core genes from each of these enrichment plots, we utilized Venn diagrams to find the overlap between cell identity genes that changed with normal aging and in the exposure signatures. In males exposed to BPA.hi, TBT, and TCDD, for cell identity genes of both parenchymal and non-parenchymal cells, we consistently observed an anti-correlation between core genes in the aging and exposure signatures, i.e attenuated aging. Specifically, for hepatocytes, core identity genes that typically increase with age showed significant overlap with core genes decreased in the exposure signatures—reflecting attenuated aging—in adult males exposed to BPA.hi, TBT, and TCDD (Figure 6c). This attenuated aging effect was similarly observed in females exposed to TBT and TCDD (Supplemental Figure 6d). Conversely, in non-parenchymal cells, we found substantial overlap in core cell identity genes of Stellate, Kupffer, NK, and endothelial cells that decrease with age and those that increased in response to TBT, TCDD, and BPA exposures (Figure 6c) and in females exposed to TBT, TCDD, and PM2.5-Chi (Supplemental Figure 6d).

In males for example, 206 core hepatocyte aging DEGs typically increased between weaning and young adulthood (normalized enrichment score [NES] = 3.93), and among these, expression was attenuated for 122 (59%; NES =-3.52), 116 (56%; NES =-4.55), and 95 (46%; NES =-2.39) in adult hepatocytes exposed to BPA, TBT, and TCDD, respectively. A similar attenuated aging was observed in non-parenchymal cells. In Stellate cells, 112 core genes normally decreased with age (NES = - 4.89), of which 68 (61%; NES = 4.33), 82 (73%; NES = 4.09), and 60 (54%; NES = 3.46) remained high in livers exposed to BPA, TBT, and TCDD, respectively. Similar overlaps were seen for core genes in Kupffer, NK-NKT, and endothelial cells, where genes that typically decreased with age remained elevated in adult livers of exposed males (Figure 6c) and females (Supplemental Figure 6d). Sex-specific responses were also observed. For instance, PM2.5-Chi attenuated expression of agingDEGs only in male hepatocytes, and TCDD-induced attenuated expression of agingDEGs in Stellate and endothelial cells was exclusive to male livers (compare Figures 6a and Supplemental Figure 6b).

We next utilized the genes in the intersection of aging and exposure signatures to identify functional pathways enriched in the attenuated aging genes for each cell type in males exposed to BPA.hi, TBT and TCDD (Figure 6d) and females exposed to PM2.5-Chi, TBT and TCDD (Supplemental Figure 6d) as determined using ORA (Supplemental Table 9b-c). In hepatocytes, attenuated agingDEGs predominantly enriched for metabolic pathways that typically increase with age, but which remained low in the adult liver of exposed animals. Conversely, in non-parenchymal cells, attenuated agingDEGs were enriched for pathways associated with cytoskeletal organization, extracellular matrix production, and immune signaling, which rather than decreasing with age, remained high in exposed animals. This inverse relationship, wherein aging processes and cell-type specificity shape polarized T2C exposure signatures, is depicted schematically in Figure 6d, highlighting the dynamic interplay between epigenomic reprogramming and cell-type-specific aging trajectories, and the systemic impact of these early-life exposures.

To confirm that the observed attenuated aging phenotype – characterized by increased or decreased agingDEG expression relative to age-matched controls – was due to disruption of the transcriptome rather than shifts in liver cell composition, we employed MuSiC (Multi-subject Single-cell deconvolution) analysis (Supplemental Figure 6e) to assess the impact of T2C exposures on liver composition of parenchymal and non-parenchymal cells. None of the exposures induced significant changes in the proportions of these cell types, with the exception for PM2.5-Chi, where a slight, but statistically significant (p < 0.003), increase in the proportion of hepatocytes occurred (from 0.995 to 1.0 and 0.994 to 0.996). Notably, this minor change in cell composition was opposite to the transcriptional reprogramming observed where hepatocytes (despite their increase in number) showed a reduction in metabolic gene expression. This confirms that the attenuated aging phenotype was driven by exposure-related transcriptional reprogramming rather than any change in cellular proportions.

We also asked to what extent the attenuated aging induced by multiple exposures was due to reprogramming of the same, or different genes, in these functional pathways. As shown in Supplemental Figure 6f using heatmaps of core identity genes and the Venn diagrams in Figure 6e, we identified agingDEGs reprogrammed by one or more exposures in hepatocytes or Stellate cells (see Supplemental Table 9). The picture that emerged for BPA.hi, TCDD, and TBT in hepatocytes was that only about a quarter (41/178) of the hepatocyte attenuated aging genes reprogrammed in the adult male liver were targeted by all three exposures. Similarly in Stellate cells, only about a third (33/102) attenuated aging genes were targeted by all three exposures. This suggests that the underlying feature shared by pathways targeted for attenuated aging – such as metabolism in hepatocytes and extracellular matrix in Stellate cells – was the plasticity of genes in these pathways to change expression with age, creating a vulnerability to early-life exposures – exposures with very different mechanisms of action, epigenetic exposure signatures, and acting on largely different gene targets within the same pathway.

### Attenuated Aging Distinguishes Healthy from Diseased Liver in Patient Populations

Non-alcoholic fatty liver disease (NAFLD) has been linked to environmental exposures, as highlighted by Foulds ^30^, and is a key precursor in the progression to hepatocellular carcinoma (HCC) ^31,32^. Among the exposures studied within T2C, TBT is specifically associated with NAFLD in male mice ^33,34^ and, with variable penetrance, the development of liver tumors ^22,23^. Epigenomic and transcriptional profiling across multiple individual mice provided an opportunity to capture heterogeneity in response, and the profiling of individual mice, rather than pooled samples, was one of the strengths of the T2C studies. For example, in the cohort of T2C C57Bl/6 male mice exposed to TBT followed out to 10 months, when uninvolved liver tissue from tumor-bearing mice was compared to age-matched identically exposed mice that did not develop tumors, distinct epigenomic and transcriptomic signatures in the two groups emerged. In TBT-exposed mice that developed tumors, the uninvolved liver tissue exhibited a signature markedly different from age-matched vehicle controls, but with many similarities to the tumors (i.e a “high-risk” TBT exposure signature), whereas liver tissue of TBT-exposed mice without tumors displayed a signature that had remained very similar to vehicle controls (i.e a “low-risk” TBT exposure signature) ^23^. Thus, while liver tumor development among male mice equivalently exposed to TBT exhibited variable penetrance, individual risk for tumor development was linked to epigenomic and transcriptomic reprogramming of the liver induced by this EDC ^23^.

If the observed heterogeneity in the epigenomic and transcriptomic responses to TBT exposure precedes tumor development, then aggregating data from individual mice (as done for TBT and other T2C exposures above) could obscure critical features of these early-life exposure effects. Indeed, when analyzed individually, young adult TBT-exposed male mice stratified into two distinct endotypes at 5 months: Endotype 1, characterized by significant transcriptomic and epigenomic alterations resembling the high-risk signature observed in liver tissue at 10 months, and Endotype 2, exhibiting a low-risk, vehicle-like signature. These distinct endotypes are illustrated in the heatmap and PCA plots in Figure 7a–b. Notably, transcriptional profiling of blood confirmed TBT exposure occurred in both Endotype 1 and Endotype 2 groups, consistent with the systemic nature of the maternal exposure paradigm.

**Figure 7.**
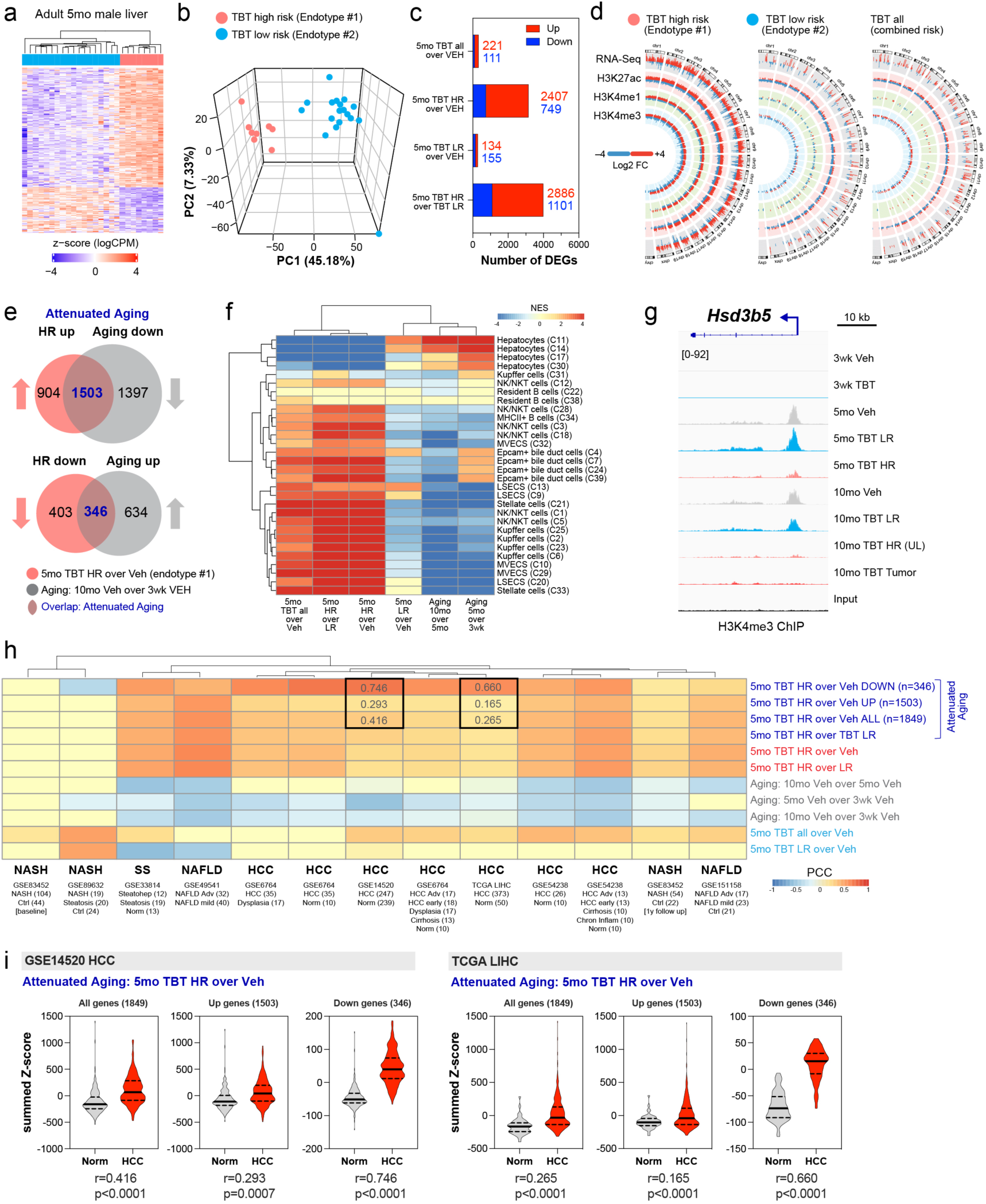
EDC reprogrammed attenuated aging epigenomic and transcriptomic signature distinguishes healthy from diseased livers. a. Heatmap and b. PCA of Endotype 1 (high risk/HR) and Endotype 2 (low risk/LR) in 5mo male mice after TBT exposure. c. Summary of exposure DEGs in HR and LR. d. Circos plots showing direction and magnitude of exposure DEPs in the HR and LR endotypes compared to age matched vehicle controls, contrasted to combined risk. e. The HR Endotype 1 exposure DEG shows attenuated aging. f. GSEA enrichment of liver cell type specific markers in 5mo and 10mo exposed animals confirms attenuated aging associated with high risk. g. IGV plot showing Endotype-specific DEPs at the *Hsd3b5* locus. h. Pearson Correlation Coefficient analysis of exposure and aging DEGs against multiple human cohorts of liver disease (p<0.05). i. Summed z-scores distribution for the HR endotype attenuated aging DEGs for two human liver cancer transcriptomic datasets (p<0.05).

Comparative analyses of the liver transcriptomes and epigenomes of Endotype 1 and Endotype 2 mice, relative to age-matched vehicle controls, revealed striking differences (Figure 7c–d and Supplemental Table 10). Endotype 1 livers exhibited a markedly distinct transcriptional profile, with 2,407 upregulated and 749 downregulated DEGs relative to vehicle controls. In contrast, Endotype 2 livers more closely resembled vehicle controls, with only 134 upregulated and 155 downregulated DEGs. This pronounced divergence highlights the robust signature of Endotype 1 transcriptomes compared to aggregated data from multiple 5-month-old animals (Figure 7c; compare with Figure 2a), and underscore a profound molecular heterogeneity in TBT response, revealing a high-risk signature that emerges prior to tumor development and demonstrating the critical value of individual-level analyses.

Accounting for this heterogeneity also better resolved the attenuated aging transcriptomic signature induced by TBT (compare Figure 7e with Figure 3d). In Endotype 1 livers, 62% (1,503/2,407) of upregulated signature genes were agingDEGs that typically decline with age, while 46% (346/749) of downregulated signature genes were agingDEGs that normally increase with age. The epigenomic profiles of Endotype 1 and Endotype 2 livers were similarly distinct, as shown in the Circos plots in Figure 7d. Endotype 1 livers exhibited significant epigenomic divergence from age-matched vehicle controls, whereas Endotype 2 livers displayed minimal changes relative to controls. Importantly, when epigenomic profiles from all 5-month TBT-exposed mice were aggregated, the overall response appeared muted, masking the pronounced changes that had occurred in mice of Endotype 1. As shown in Figure 7d, Endotype 1 livers exhibited substantially greater gains and losses in H3K27ac, H3K4me1, and H3K4me3 compared to Endotype 2 livers, further highlighting the critical differences between these two response types.

GSEA for parenchymal and nonparenchymal cell identity genes, as shown in the heat map in Figure 7f, revealed the polarized attenuated aging phenotype seen with aggregated data was clearly driven by reprogramming in the Endotype 1 livers, as detailed in Supplemental Table 11. Across parenchymal and nonparenchymal cell types, Endotype 1 with “high-risk” transcriptomic and epigenomic features, whether compared to vehicle or Endotype 2 “low-risk” livers, formed a distinct cluster, and had a more robust anti-correlation with cell-type identity genes-characteristic of attenuated aging-than was seen with the aggregated transcriptomes of all 5 month mice (Figure 7f). This was the case for both the negative enrichment of hepatocyte cell identity genes that normally increase with age, as well as the positive enrichment for nonparenchymal cell genes that normally decrease with age. In contrast, the Endotype 2 low-risk livers clustered separately, and were aligned directionally and with cell specificity, with normal aging signatures (Figure 7f). An example of the epigenomic correlates for this reprogramming is shown in Figure 7g for the hepatocyte metabolic gene hydroxy-delta-5-steroid dehydrogenase, 3 beta-and steroid delta-isomerase 5 (*Hsd3b5*). From weaning to early (5 months) and later (10 months) adulthood, *Hsd3b5* expression in hepatocytes normally increases 16-to 64-fold, accompanied by gain of H3K4me3 at the *Hsd3b5* promoter (Figure 7g). The gain of H3K4me3 seen in vehicle controls with age also occurs in Endotype 2 TBT-exposed livers at 5 months, and in low-risk 10-month TBT-exposed livers of animals that did not develop tumors. However, this H3K4me3 DEP was absent in livers of 5-month-old Endotype 1 and uninvolved liver tissue of 10-month-old high-risk mice that developed liver tumors. In these tissues, *Hsd3b5* expression was ∼100-fold less than what is seen in age-matched controls.

To establish the translational relevance of polarized attenuated aging in the liver, we next performed Pearson’s correlation analyses using transcriptomic data from multiple clinical liver diseases: NAFLD, Non-Alcoholic Steatohepatitis (NASH), simple steatosis (SS), and hepatocellular carcinoma (HCC) patient cohorts. These liver transcriptomes were compared to transcriptomic signatures of normal liver aging obtained in mice from weaning to young adulthood and young to later adulthood, Endotype 1 high-risk and Endotype 2 low-risk TBT exposure 5 month signatures. Strikingly, transcriptional signatures from diseased human livers and HCC showed a similar negative correlation with genes that normally change during liver aging (i.e., attenuated aging) and a strong positive correlation with the Endotype 1, high-risk attenuated aging signature (Figure 7h).

This correlation was strongest between human HCC cohorts and TBT signature genes that were downregulated (i.e. enriched for hepatocyte identity genes). This is illustrated using violin plots over gene expression data from tumor and adjacent non-tumor tissue in HCC patients (GEO GSE14520 and TCGA) (Figure 7i). In both datasets, attenuated agingDEGs from the high-risk Endotype 1 TBT exposure group that were *decreased* (i.e. enriched for hepatocyte-specific genes) most effectively distinguished human HCC from normal livers. In the GEO GSE14520 dataset, attenuated agingDEGs that *decreased* in the Endotype 1 high-risk signature showed a robust positive correlation with HCC (r = 0.746), compared to genes that were *increased* (i.e. enriched for nonparenchymal cell genes) (r = 0.293) or when all genes were aggregated (r = 0.416) (Figure 7i left). The TCGA dataset similarly confirmed this association: attenuated agingDEGs that *decreased* in the high-risk TBT signature better differentiated normal liver from HCC (r = 0.660) than genes that were increased (r = 0.165) or all genes aggregated (r = 0.265) (Figure 7i right). These findings underscore a fundamental link between attenuated expression of agingDEGs and liver disease, including HCC, a linkage conserved across species, and vulnerable to disruption by early-life environmental exposures.

## DISCUSSION

The NIEHS TaRGET II Consortium has undertaken the most comprehensive parallel transcriptomic and epigenomic profiling effort to date to shed light on the persistent effects of early-life environmental exposures on the liver. These studies revealed that epigenetic histone modifications – particularly those associated with enhancers – and age-related gene expression changes are key targets for reprogramming in response to diverse early-life environmental insults. The findings suggest that the inherent plasticity required for programmed epigenomic aging, along with shifts in the trajectory of aging-associated gene expression, underpins the liver’s vulnerability to reprogramming by environmental exposures. Crucially, aberrant expression of agingDEGs, particularly the attenuation of their normal aging trajectory, was linked to risk for development of liver tumors in mice and correlated with development of liver disease and HCC in patient cohorts. Together, these findings provide groundbreaking insights into how early-life exposures disrupt the epigenome and transcriptome, while also establishing environmental exposures as a powerful lens through which to unravel fundamental mechanisms driving human disease.

## MATERIALS AND METHODS

### Animal Exposure Paradigm and Tissue Collection

C57BL/6 (B6; Jackson Laboratory, Bar Harbor, ME) mice were used for these experiments with the exception of the lead (Pb) and phthalate (DEHP) exposures in which wild-type non-agouti *a/a* mice from an over 230-generation colony of viable yellow agouti (*A^vy^*) mice, which are genetically invariant and 93% identical to the C57BL/6J strain were used ^35^. Mice were exposed to toxicants perinatally through maternal dietary, water consumption, inhalation or oral gavage with an exposure period that spanned pre-conception to weaning; Supplemental Figure 1a contains a schematic describing the experimental design of the mouse perinatal exposure study. Two weeks prior to mating virgin female dams (6-8 weeks old) were randomly assigned to one of nine exposure groups: 1) 50 mg/kg diet of BPA (BPA high); 2) 50 µg/kg diet of BPA (BPA low); 3) 5 mg DEHP/kg diet/day; 4) 32 ppm Pb-acetate drinking water; 5) PM 2.5 Baltimore; 6) PM 2.5 Chicago; 7) tributyltin (TBT); 8) 1 ug/kg dioxin (TCDD); or 9) control. Dams were mated with virgin males (7-9 weeks old) two weeks being placed on experimental diets or water, and exposure continued through pregnancy and offspring weaning at postnatal day 21 (PND21). Animals were housed in polycarbonate-free cages, and remained on a 12- hour light/dark cycle. All procedures were approved by the Institutional Animal Care and Use Committees affiliated with each production site and conducted in accordance with experimental procedures outlined by the NIEHS TaRGET II Consortium and the highest animal welfare standards ^9^. For these experiments, the litter was considered as the N, and only one male and female mouse from each litter contributed data to any one timepoint.

At 3 weeks (PND21), all offspring were weaned and moved to control water and control chow. Consortium-wide, one male and one female mouse per litter/exposure group was euthanized at 3 weeks of age for tissue collection and molecular analyses, a second set of one male and one female were maintained until 5 months of age, and a third set of one male and one female were maintained until 10 months of age. This paper evaluates livers obtained from male and females animals collected at all three time points. For organ and tissue collection, mice were fasted during the light cycle for four hours beginning in the morning, and then underwent CO_2_ euthanasia and cardiac puncture in the afternoon, followed by whole-body perfusion with saline (Sigma), followed by dissection of the liver, which was immediately flash frozen in liquid nitrogen and stored at-80°C. Tissue collection and processing followed protocols developed by the TaRGET II Methods Working Group, using standardized operating procedures, which can be found located on our website https://targetepigenomics.org/documents/, including SOP_Blood_Processing, SOP Mouse_Liver_Collection, and SOP_Whole_Mouse_Perfusion. For each mouse, one investigator administered the treatment and was therefore aware of the treatment group allocation; all investigators completing subsequent molecular assays were blinded to treatment group, until treatment group was analyzed during bioinformatic analyses. Numbers of mice in each arm used for the liver analysis in this study are shown in Table 4 below.

**Table 4.**
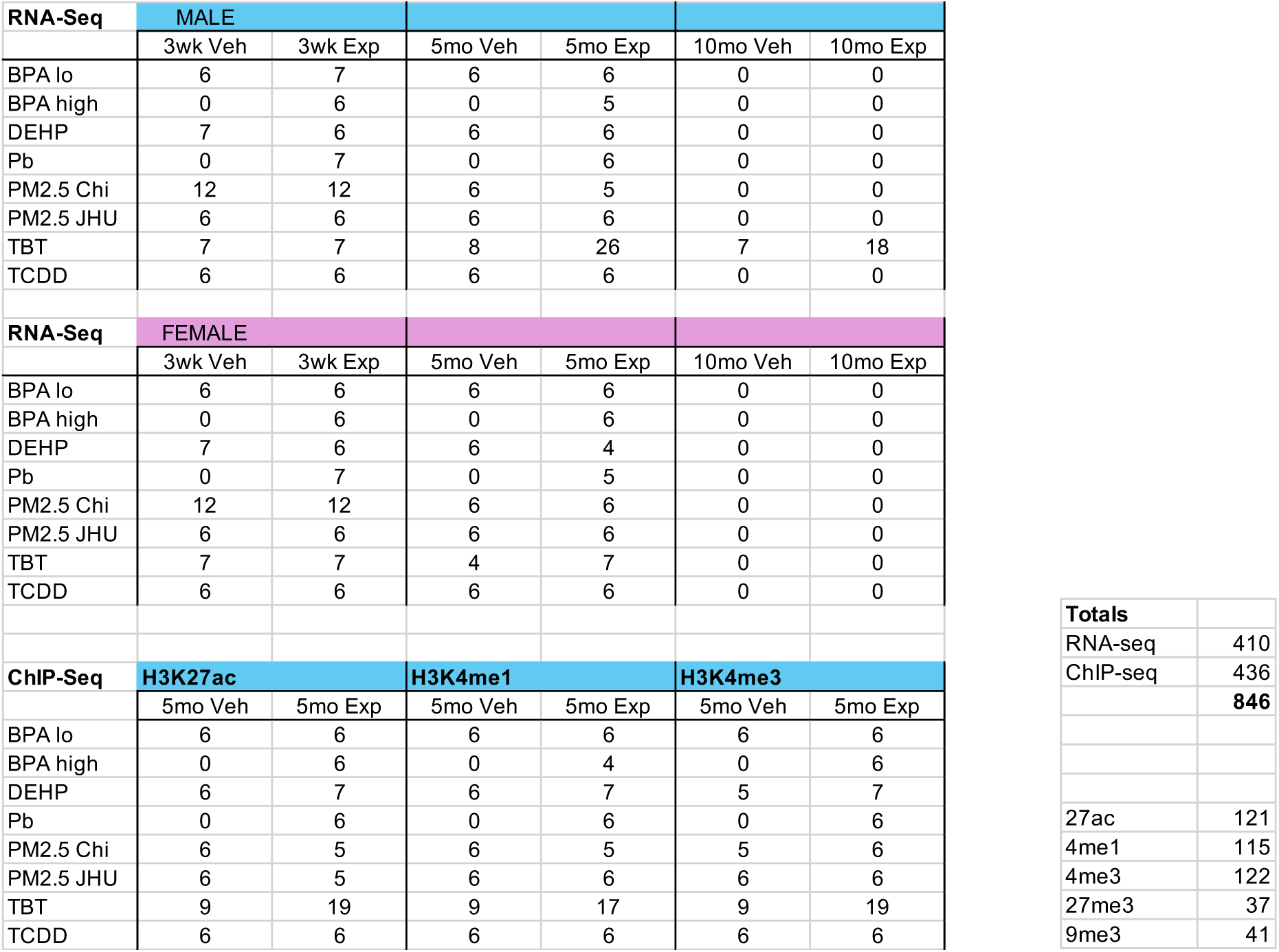
Number of animals per omic.

| RNA-Seq | MALE |  |  |  |  |  |
| --- | --- | --- | --- | --- | --- | --- |
|  | 3wk Veh | 3wk Exp | 5mo Veh | 5mo Exp | 10mo Veh | 10mo Exp |
| BPA lo | 6 | 7 | 6 | 6 | 0 | 0 |
| BPA high | 0 | 6 | 0 | 5 | 0 | 0 |
| DEHP | 7 | 6 | 6 | 6 | 0 | 0 |
| Pb | 0 | 7 | 0 | 6 | 0 | 0 |
| PM2.5 Chi | 12 | 12 | 6 | 5 | 0 | 0 |
| PM2.5 JHU | 6 | 6 | 6 | 6 | 0 | 0 |
| TBT | 7 | 7 | 8 | 26 | 7 | 18 |
| TCDD | 6 | 6 | 6 | 6 | 0 | 0 |
| RNA-Seq | FEMALE |  |  |  |  |  |
|  | 3wk Veh | 3wk Exp | 5mo Veh | 5mo Exp | 10mo Veh | 10mo Exp |
| BPA lo | 6 | 6 | 6 | 6 | 0 | 0 |
| BPA high | 0 | 6 | 0 | 6 | 0 | 0 |
| DEHP | 7 | 6 | 6 | 4 | 0 | 0 |
| Pb | 0 | 7 | 0 | 5 | 0 | 0 |
| PM2.5 Chi | 12 | 12 | 6 | 6 | 0 | 0 |
| PM2.5 JHU | 6 | 6 | 6 | 6 | 0 | 0 |
| TBT | 7 | 7 | 4 | 7 | 0 | 0 |
| TCDD | 6 | 6 | 6 | 6 | 0 | 0 |
| ChIP-Seq | H3K27ac |  | H3K4me1 |  | H3K4me3 |  |
|  | 5mo Veh | 5mo Exp | 5mo Veh | 5mo Exp | 5mo Veh | 5mo Exp |
| BPA lo | 6 | 6 | 6 | 6 | 6 | 6 |
| BPA high | 0 | 6 | 0 | 4 | 0 | 6 |
| DEHP | 6 | 7 | 6 | 7 | 5 | 7 |
| Pb | 0 | 6 | 0 | 6 | 0 | 6 |
| PM2.5 Chi | 6 | 5 | 6 | 5 | 5 | 6 |
| PM2.5 JHU | 6 | 5 | 6 | 6 | 6 | 6 |
| TBT | 9 | 19 | 9 | 17 | 9 | 19 |
| TCDD | 6 | 6 | 6 | 6 | 6 | 6 |

|  |  |
| --- | --- |
| **Totals** |  |
| RNA-seq | 410 |
| ChIP-seq | 436 |
|  | **846** |
| 27ac | 121 |
| 4me1 | 115 |
| 4me3 | 122 |
| 27me3 | 37 |
| 9me3 | 41 |

### RNA-Seq Analysis

RNA-seq was performed as described previously ^22,23^. Briefly, RNA isolated from frozen tissue and treated with DNaseI was quality verified using a Bioanalyzer. Library preparation and RNA-seq analysis was conducted in one of two places: The MD Anderson Science Park Next Generation Sequencing Core facility using the Illumina TruSeq Stranded Total RNA Protocol and the Hiseq 3000 SBS platform or The University of Houston Core facility using the QIAseq Stranded Total RNA Library Kit and the NextSeq 500 platform. Paired-end reads were trimmed using TrimGalore (https://doi.org/10.5281/zenodo.7598955) and mapped using HISAT2 ^36^ to the mouse genome build mm10, then gene expression data was quantified using featureCounts ^37^ and the GENCODE gene model ^38^. Differentially expressed genes (DEGs) were identified using the R package EdgeR ^39^, with significance achieved for a false discovery rate (FDR) adjusted p-value lower than 0.05 and a fold change exceeding 1.5x. Principal component analysis (PCA), volcano plots, and heatmaps visualizations were generated using the R statistical system. UpSet plots were generated using the R ComplexHeatmap package ^40^. Cell type deconvolution was performed using the MuSiC R package ^41^ using a single-cell liver reference ^29^.

Over-representation analysis (ORA) was performed using DEGs to detect enrichment of gene sets corresponding to pathways and biological processes using publicly available gene signatures. Following the Molecular Signature Database methodology (MSigDB), a hypergeometric test was used to assess the enrichment, with significance achieved at FDR-adjusted p-value < 0.05. Gene Set Enrichment Analysis (GSEA) was performed using the GSEA package, with significance achieved at FDR<0.25. We performed ORA and GSEA against the Gene Ontology Biological Process (GOBP), HALLMARK, KEGG, and REACTOME gene set compendia as compiled by MSigDB v7.5.1 ^42^, and against liver cell type gene signatures based on a single-cell liver reference ^29^.

Networks based on enriched pathways in distinct comparisons were derived using a Python script, with node sizes equal to the overall number of significantly enriched pathways, and edge size equal to the number of commonly enriched pathways between two different comparisons. The Cytoscape platform was used for network visualization ^43^. To determine the significance of overlap of enriched GSEA pathways, we used permutation testing with 10,000 random permutations, matching the number of significant pathways in each comparison. The results of the permutations were fitted to a normal distribution, then the final p-value was determined using the R statistical system.

### ChIP-Seq Analysis

ChIP-seq was performed as described previously ^22,23^. Briefly, chromatin was cross-linked with formaldehyde, the reaction stopped with glycine, and sonicated with a Bioruptor to obtain fragment sizes of 100–300 bp for ChIP-seq. ChIP-validated antibodies used were directed against H3K4me3 (Active Motif #39915), H3K4me1 (Abcam ab8895), H3K27ac (Abcam ab4729), H3K27me3 (Active Motif #61017), and H3K9me3 (Active Motif #61013). Immunoprecipitated DNA was recovered and purified using the QIAquick PCR Purification kit (Qiagen) and DNA concentration was measured using a NanoDrop spectrophotometer. ChIP-sequencing was performed by the University of Houston Core facility. Sequencing libraries were prepared using the QIAseq FX DNA library kit or the QIAseq Ultralow Input Library Kit and the NextSeq 500 platform. Data quality was assessed with FastQC, then mapped using BOWTIE2 ^44^ to the mouse genome build mm10; enriched regions (peaks) were called using MACS2 ^44^. Differentially expressed peaks (DEPs) were determined using diffReps ^45^, with significance achieved at FDR<0.05 and fold change exceeding 1.5x. ChIP-Seq data was visualized using the Circos platform ^46^, along with genome-wide maps using BEDTools and the Integrative Genomics Viewer (IGV; version 2.17.4). Heatmaps of ChIP-Seq signal for differentially enriched peaks (DEPs) were generated using the Deeptools software ^47^.

Quality control for ChIP-Seq was determined using the synthetic Jensen-Shannon distance for both active and repressive marks, which is based on information theory as the distance from an ideal input sample ^48^. We report the above metric for the ChIP-Seq data by aggregating samples for each histone modification and for ENCODE mouse liver ChIP-seq data (encodeproject.org) using the same marks. ChIP-seq QC data are provided in Supplemental Table 12.

Genes associated with DEPs within 3kb from transcription start sites (TSS) were determined using bedtools ^49^ software. Genes associated with DEPs at mouse enhancers were determined using bedtools and the Enhancer Atlas database (EnhAtl) ^25^. EnhAtl provides enhancer-gene chromatin loops; since we are assessing regulation across a mixture of tissues, we first used bedtools to merge enhancers from EnhAtl, thus generating enhancer anchors (Supplemental Table 4); next, we used an in-house Python code to determine enhancers that overlap with DEPs via bedtools and then associated affected enhancer anchors with their target genes.

### Signature Based Analysis Using Human Liver Transcriptomes

We used a collection of human liver transcriptomes compiled by the GTEx consortium ^50^. For each gene signature, we used Biomart to convert mouse gene symbols to human homologs. Each gene was converted to z-score across all samples. A signature score for a sample was computed by adding the z-scores of increased DEGs and subtracting the z-scores of decreased DEGs. We computed signature correlation using the Pearson Correlation Coefficient, with significance achieved for p<0.05.

We integrated these signatures with datasets from HCC samples in the Cancer Genome Atlas (TCGA) LIHC ^51^, GSE54238 ^52^, GSE6764 ^53^, and GSE14520 ^54^; NASH samples in GSE83452 ^55^ and GSE89632 ^56^; and NAFLD samples in GSE49541 ^57^ and GSE151158 ^58^. For each cohort, a gene signature activity score was derived by scaling every gene by converting it to a z-score and computing the summed z-score for all genes ^59,60^. Association of signature activity scores with liver disease was assigned by encoding the disease states with numerical values in order of severity, then computing a Pearson correlation coefficient, with significance achieved for p<0.05.

## DATA AVAILABILITY

RNA-seq data and ChIP-seq data can be accessed at the TaRGET II data portal (https://targetepigenomics.org/) and an accompanying database (http://toxitarget.com/) as well as the Gene Expression Omnibus repository (GSE146508). Additional samples were deposited to the Gene Expression Omnibus repository (accession numbers GSE272929, GSE289162, GSE272927). Custom codes are available at the Zenodo repository 10.5281/zenodo.15086523.

## Supporting information

SuppTable1

SuppTable2

SuppTable3

SuppTable4

SuppTable5

SuppTable11

SuppTable6

SuppTable7

SuppTable8

SuppTable9

SuppTable10

SuppTable12

