## Supplemental_Figures for "Early-Life Environmental Exposures Reprogram Epigenomic Aging to Alter Gene Expression Trajectories"

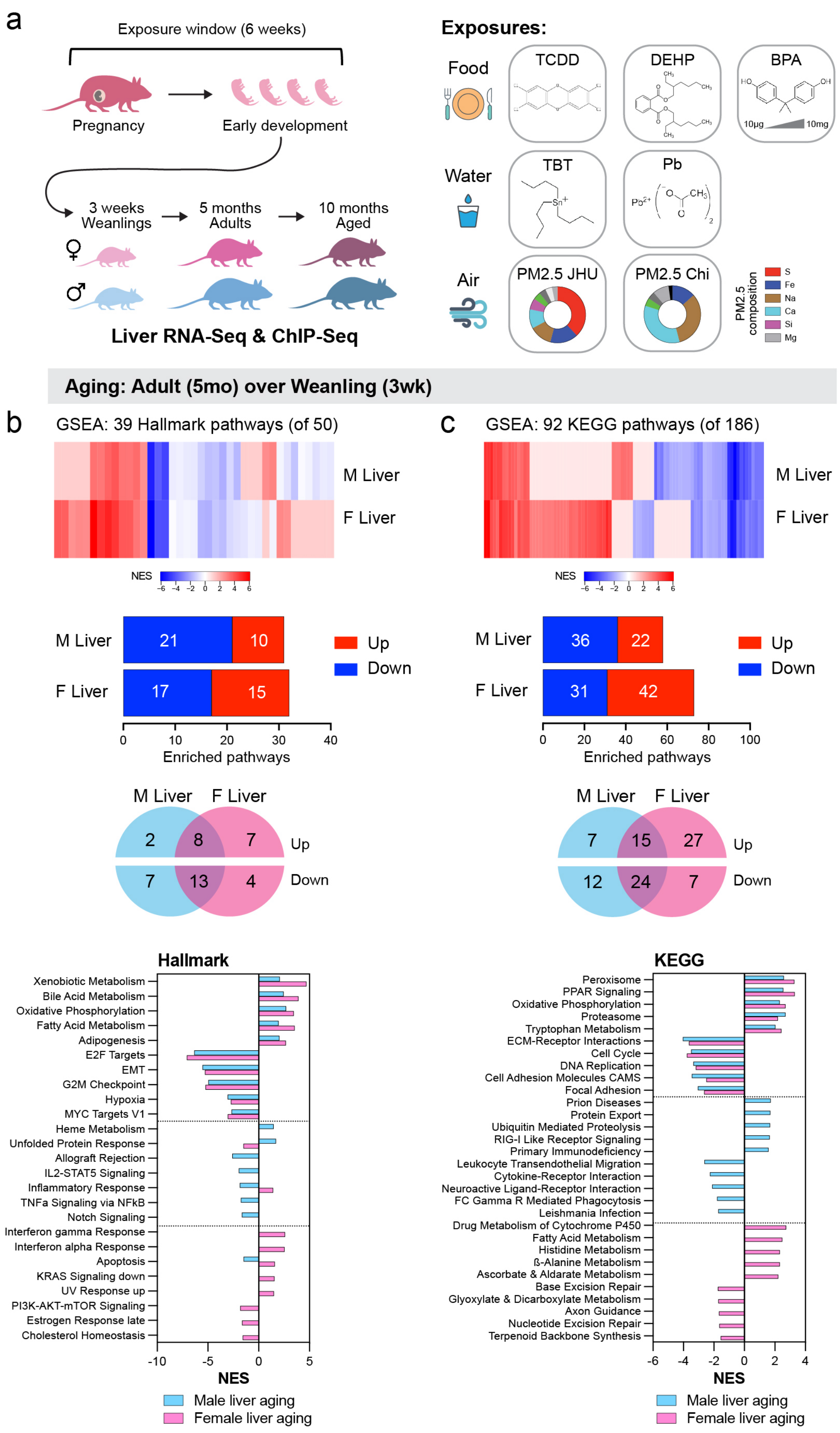


**Supp Figure 1. GSEA enriched pathways in male and female mice aging in the 3-week to 5-month time window based on multiple compendia.** a. Experimental overview. Pathway enrichment for Hallmark (b) and KEGG (c) compendia.

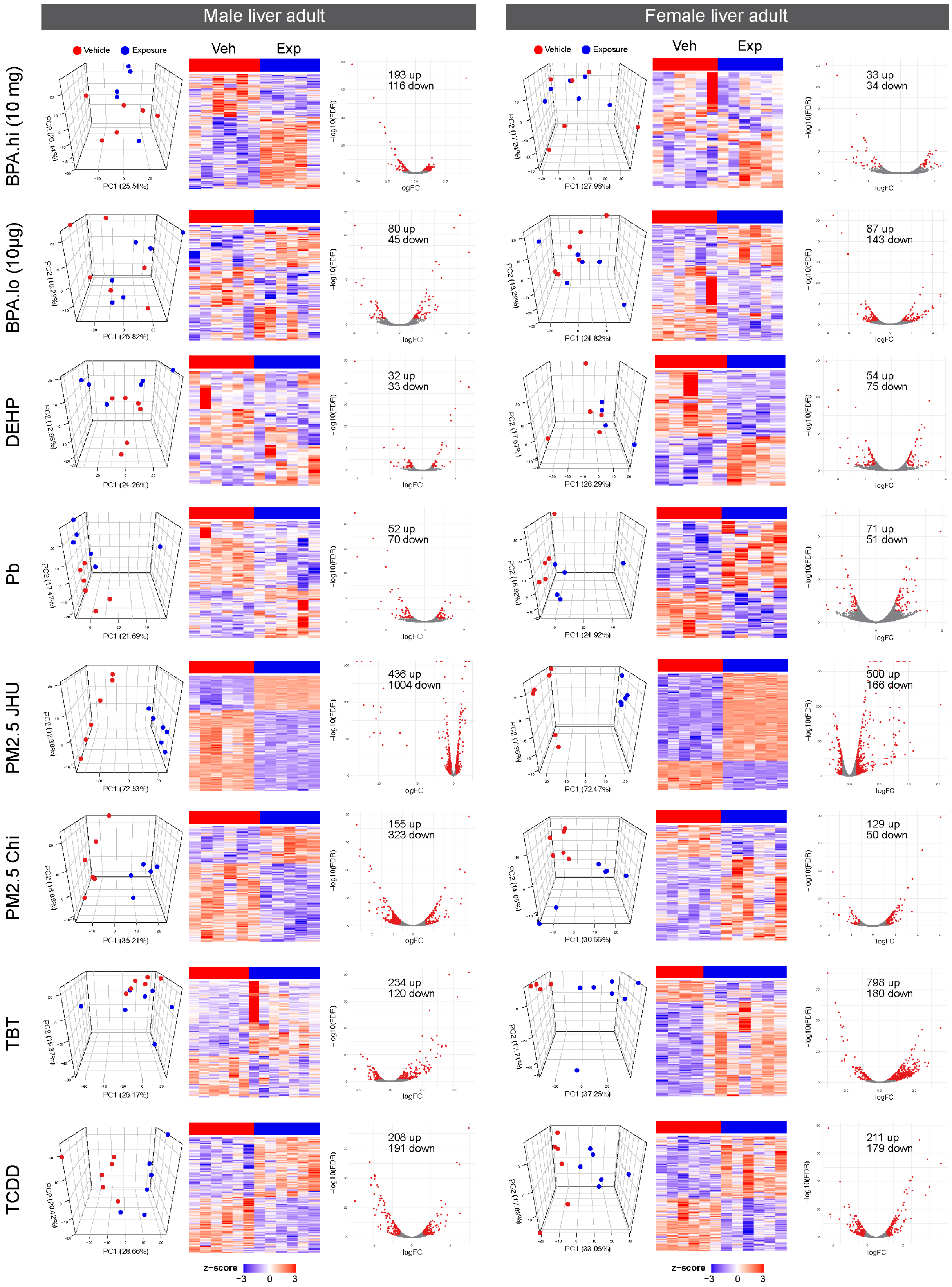


**Supp Figure 2. Transcriptome effects of early-life toxicant exposure.** For each exposure, PCA and volcano plots are shown based on all detected genes, and heatmaps based on the differentially expressed genes.

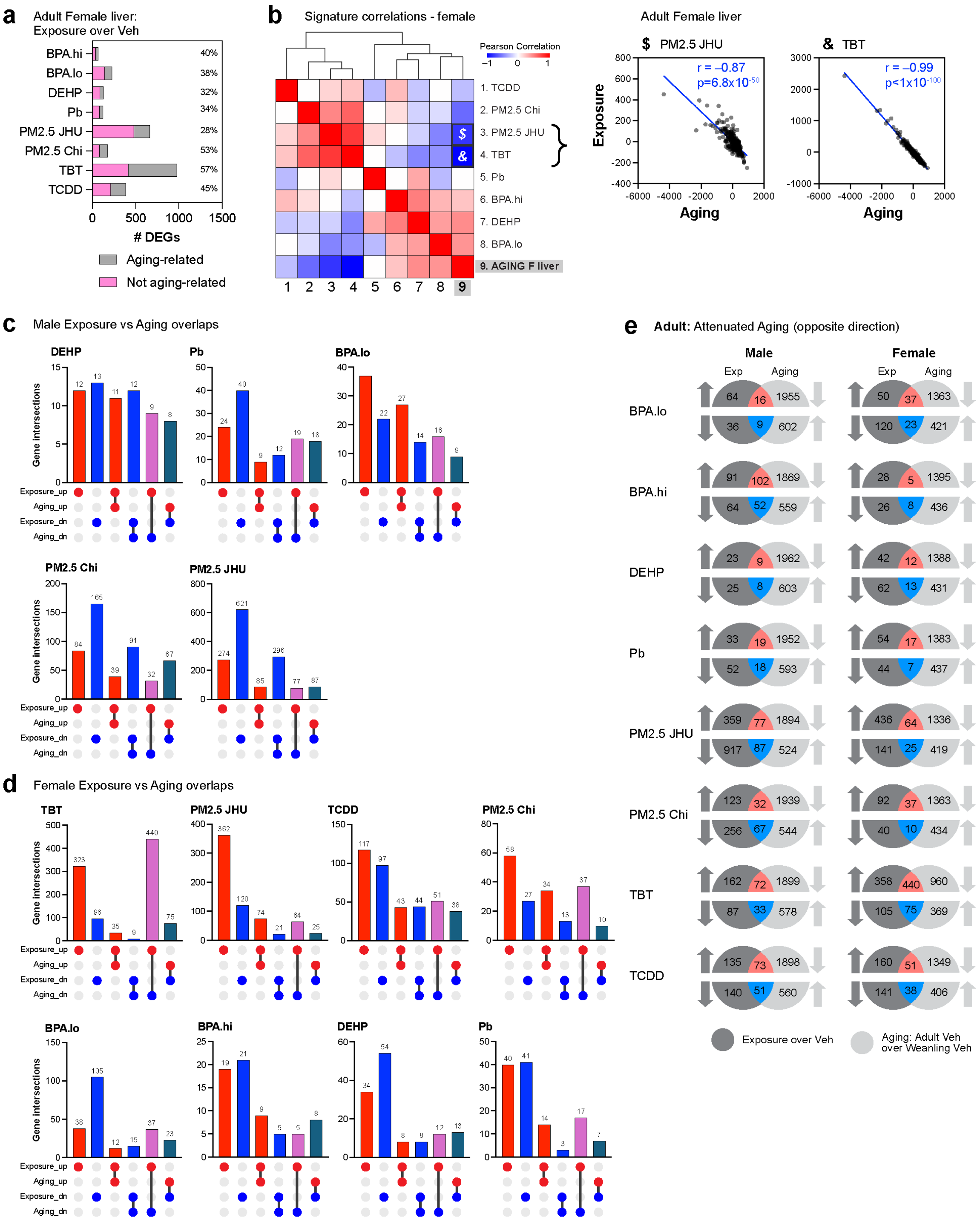


**Supp Figure 3. Overlap between exposure response and aging DEGs in both in male and female mice.** a. Bar graph summarizing overlap of aging DEGs with exposure signatures in female mice irrespective of direction. b. Matrix of Pearson's correlation coefficients (p<0.05) between summed z-scores for exposure exposure signatures and agingDEGS in female mice over the GTEx liver transcriptome dataset. c. UpSet plots showing overlap between aging DEGs and selected exposures Pb, DEHP, BPA.lo, PM2.5-JHU, and PM2.5-Chi. d. UpSet plots between agingDEGs and each exposure signature in adult female mice. e. Venn diagrams for aging and exposure DEG overlaps overlaps associated with attenuated aging in male and in female mice.

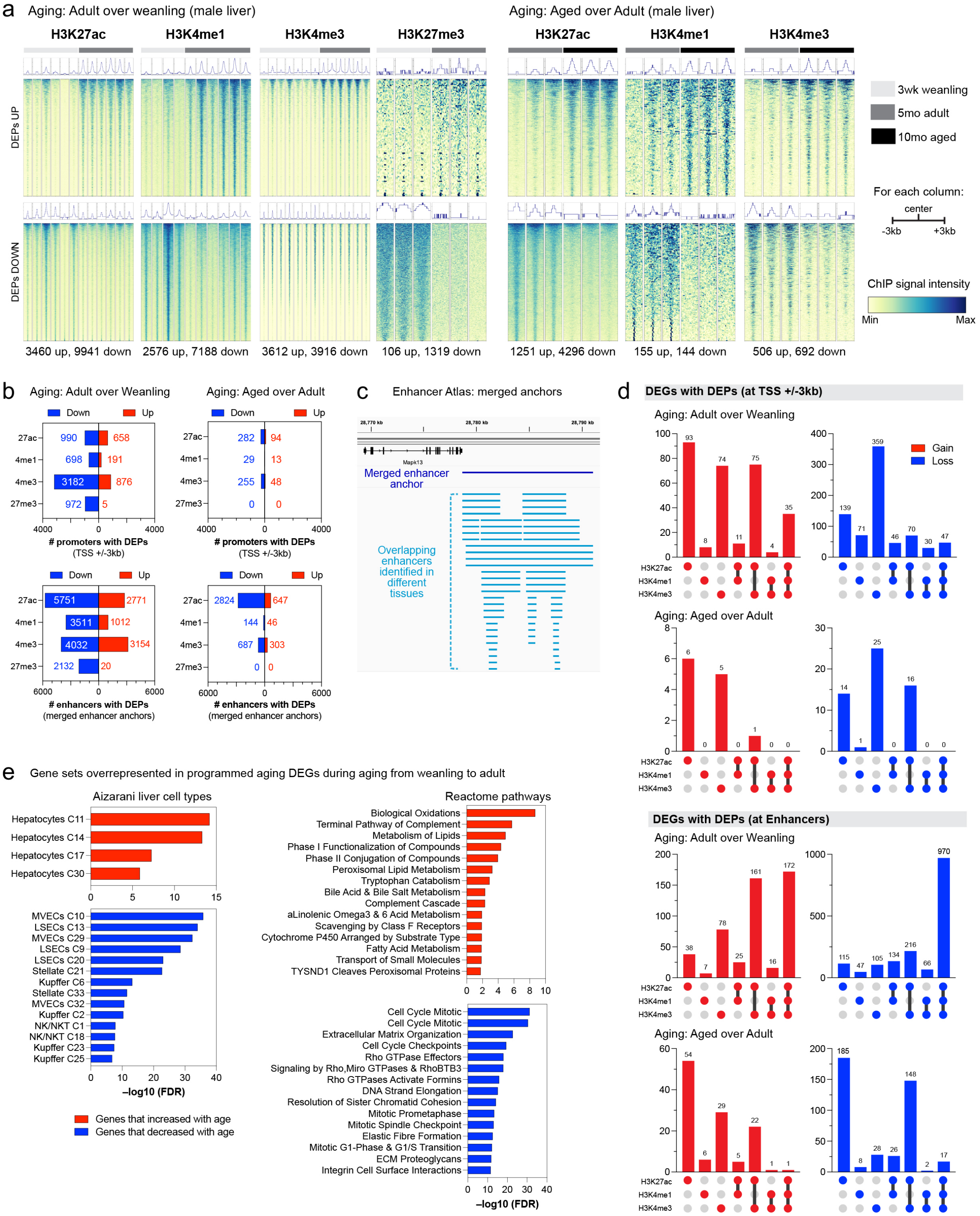


**Supp Figure 4. Epigenome programming associated with aging from weanling to young to aged adult** a. Signal heatmaps for active histone marks agingDEPs from 3 weeks to 5 months and from 5 to 10 months. b. Global changes in DEPs at gene promoters and enhancers. c. Merged enhancer anchor analysis strategy. d. Combinatorial changes of concordant agingDEP programming via TSS and enhancers for aging DEGs from 3 weeks to 5 months and from 5 months to 10 months. e. Cell type markers ORA for concordantly programmed aging DEGs using the Aizarani liver compendium or pathway ORA for concordantly programmed aging DEGs using the Reactome compendium.

### Supplemental Figure 5 (multiple pages)

a

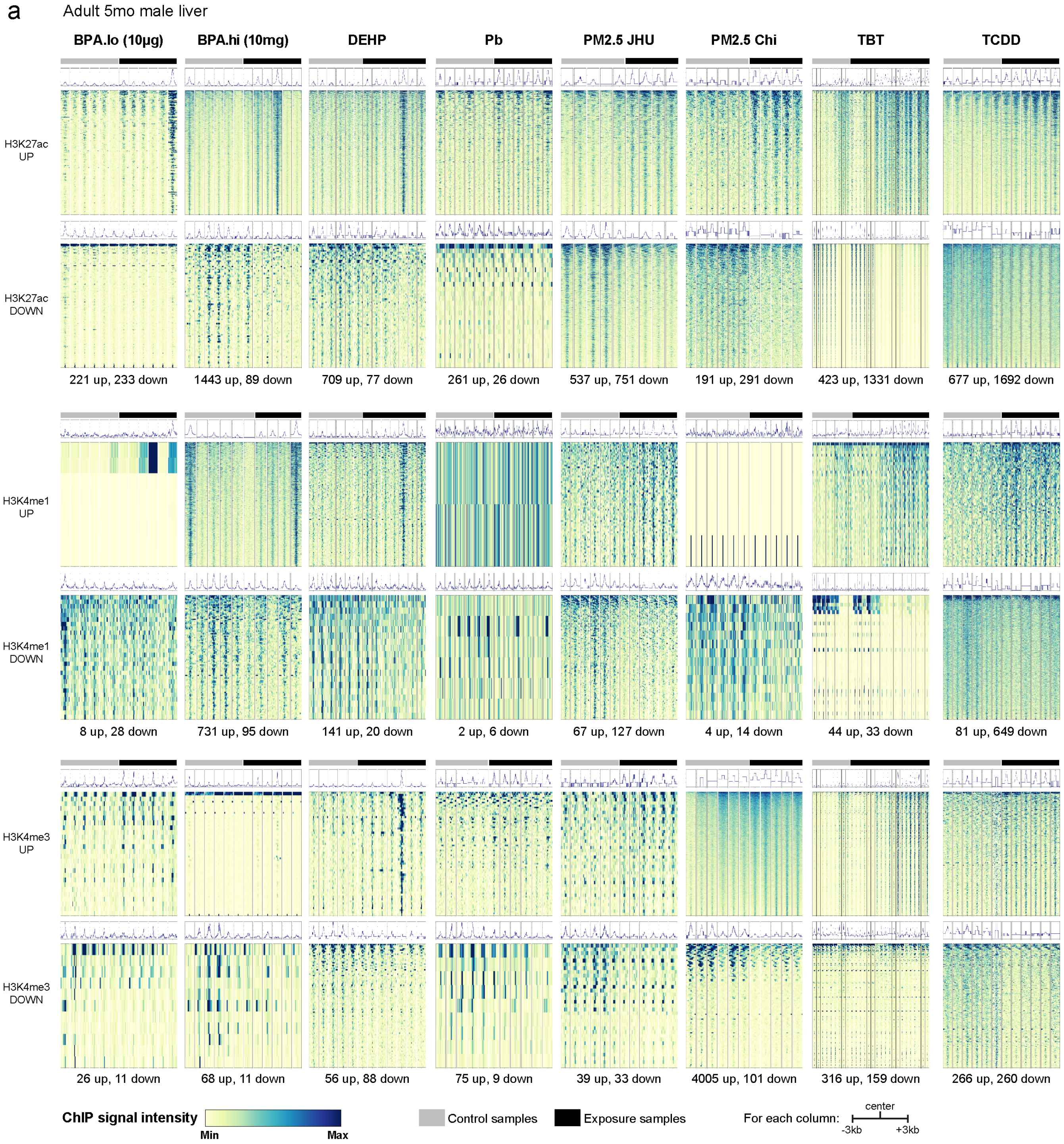

**Supplemental Figure 5 (multiple pages)**

b

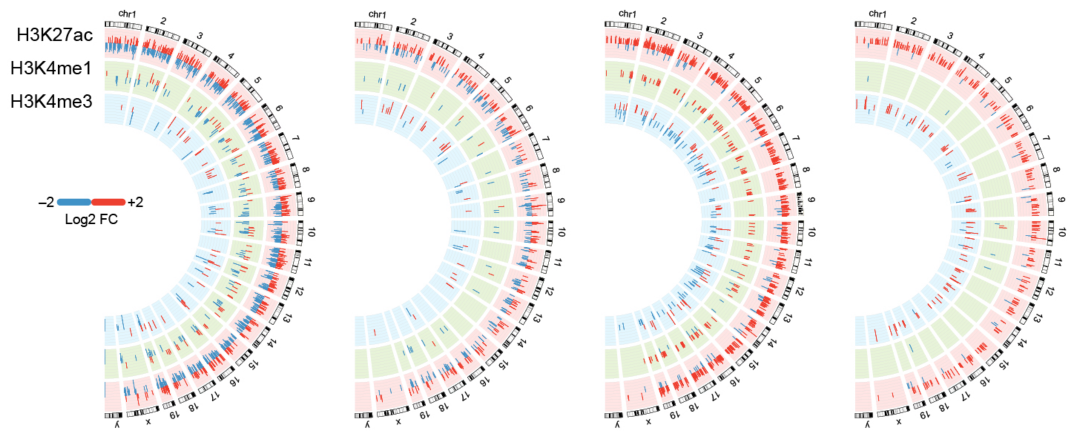


c

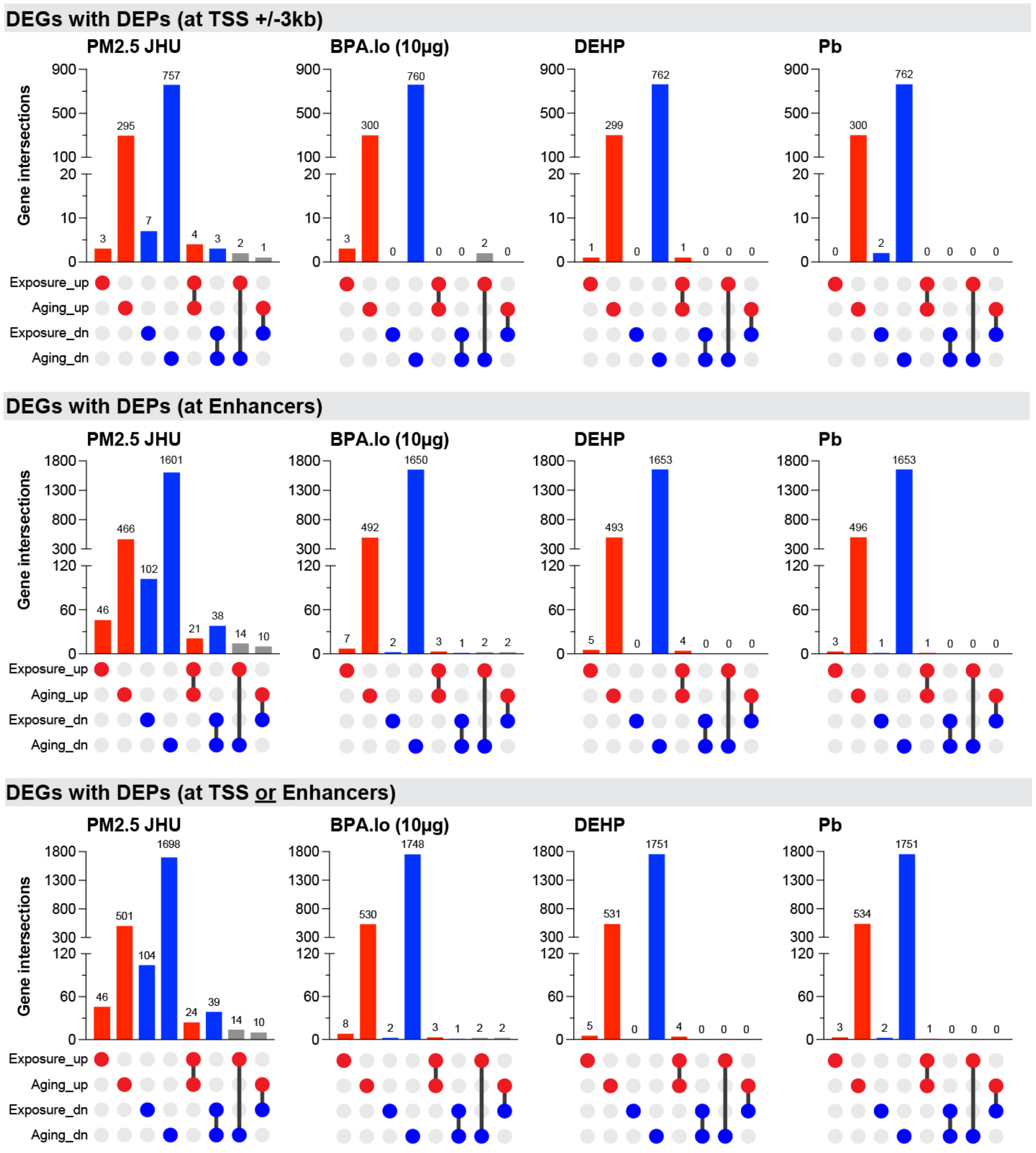

**Supplemental Figure 5 (multiple pages)**

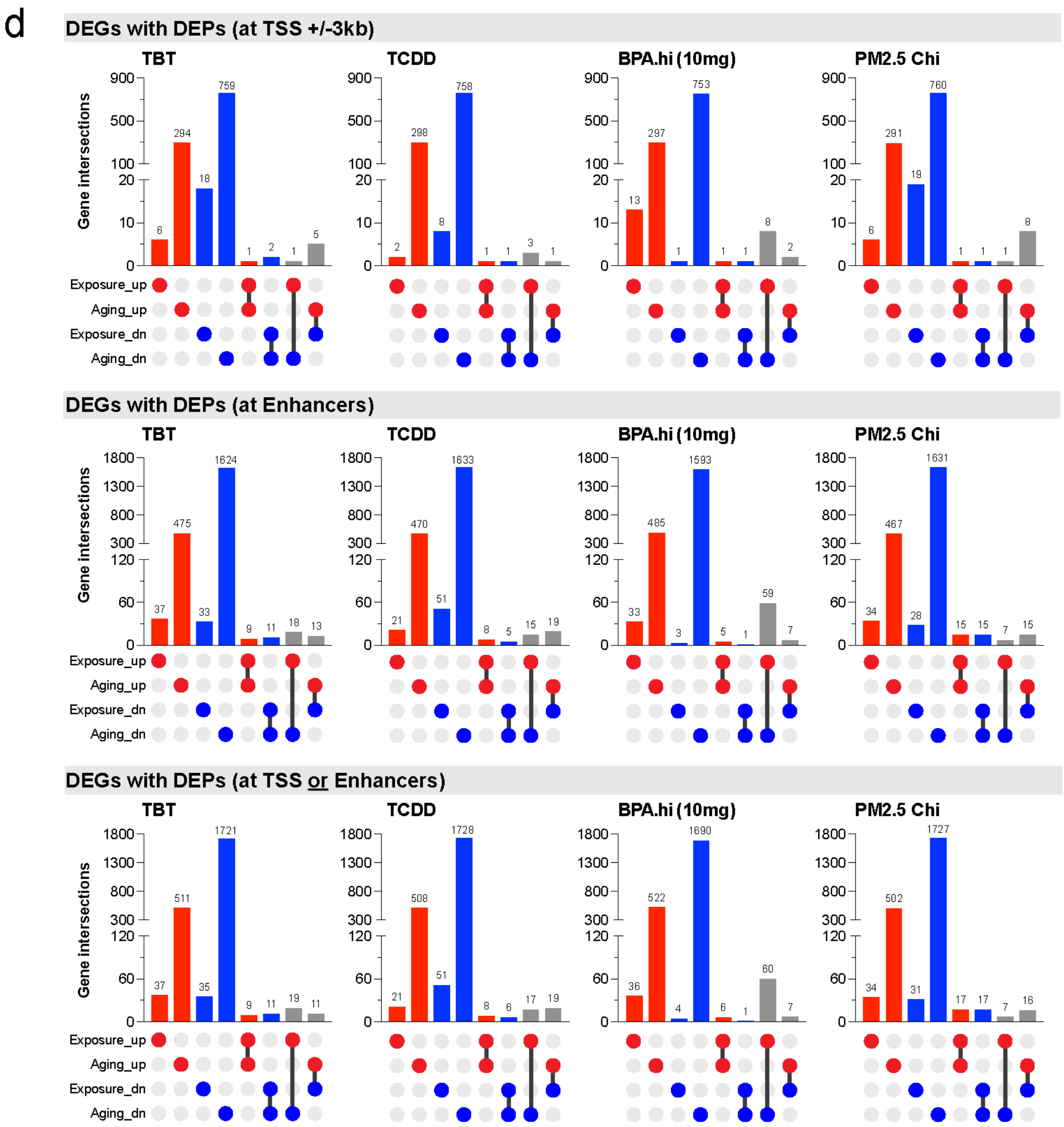

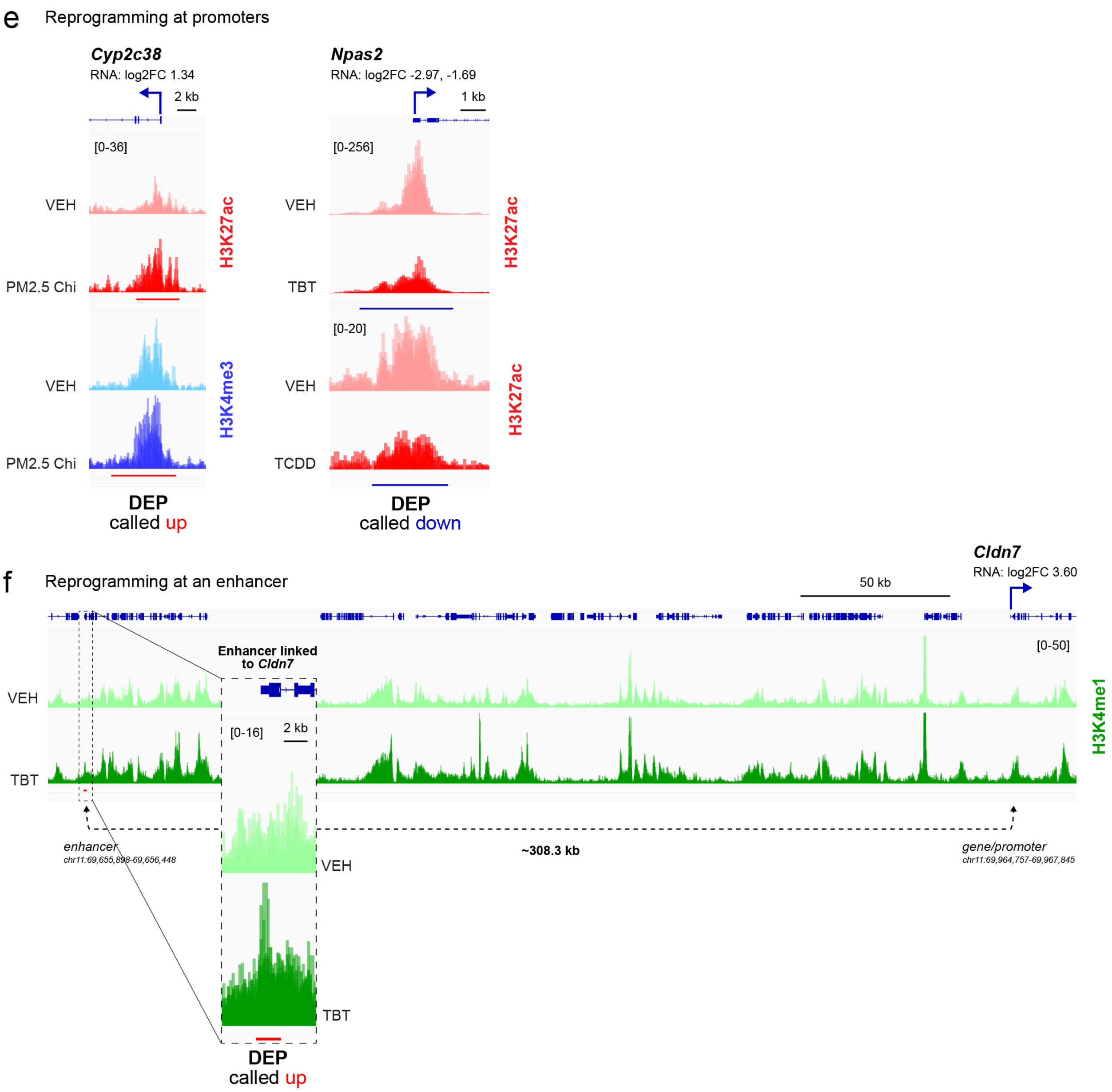


**Supp Figure 5. Epigenome reprogramming associated with early-life toxicant exposures.** a. Signal heatmaps for all active marks and exposures. b. Circos plots indicating DEP direction and magnitude of changes for PM2.5-JHU, BPA.lo, DEHP, and Pb. c. UpSet plots for overlap of aging DEGs and exposure DEGs for PM2.5-JHU, BPA.lo, DEHP, and Pb. d. UpSet plots for overlap of aging DEGs and exposure DEGs for TBT, TCDD, BPA.hi, and PM2.5-Chi. IGV examples for reprogramming at promoters (e) or an enhancer (f).

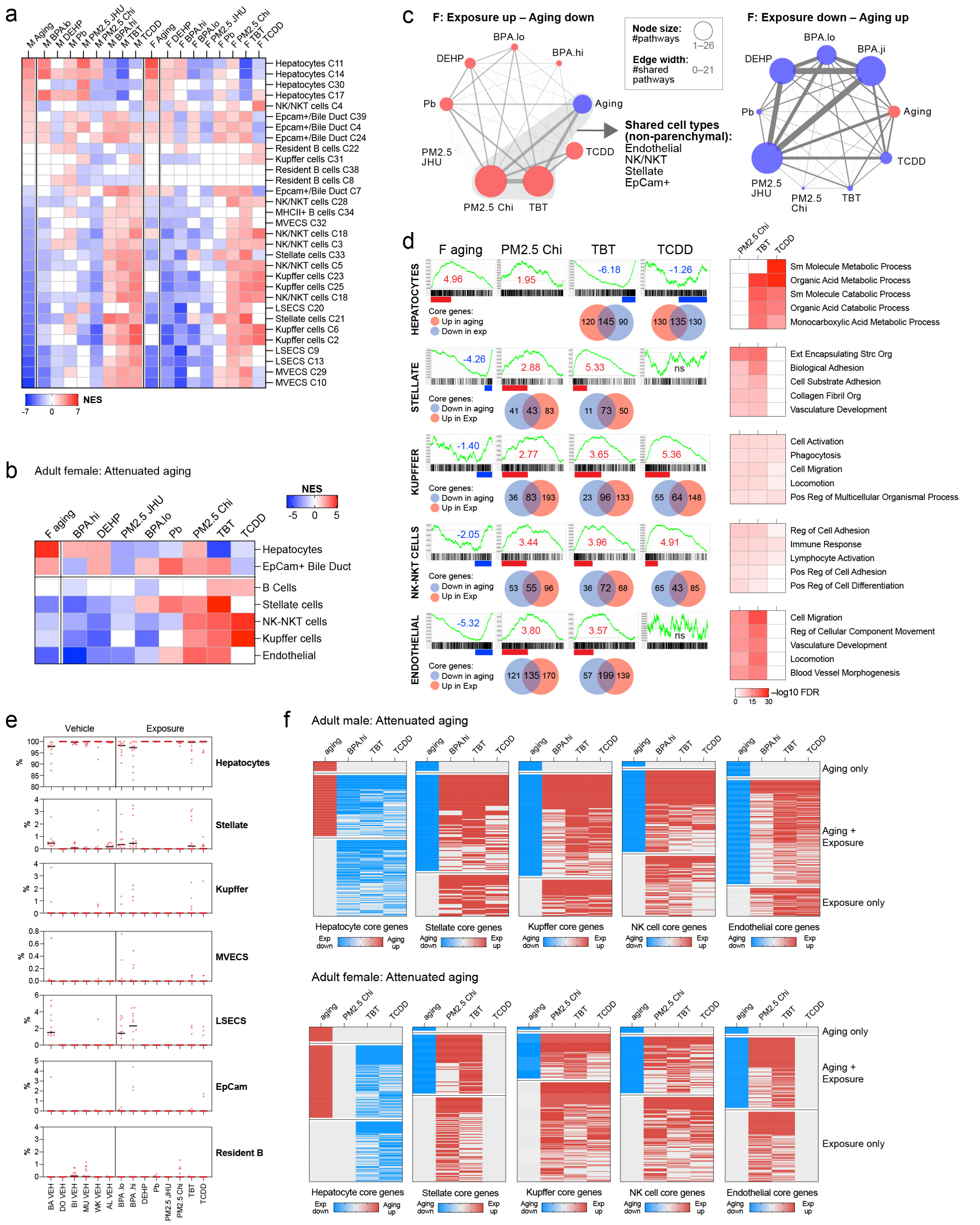


**Supp Figure 6. Environmental exposure induced cell type specific attenuated aging polarized effects.** a. GSEA enrichment heatmap for T2C exposures in males and females using Aizarani 31 cell types. b. GSEA heatmap for female 5-month using Aizarani 8 cell types. c. Network analysis of cell identity genes vs aging in females. d. GSEA enrichment plots with Venn diagrams for overlaps between aging and exposure core genes from PM2.5-Chi, TBT, and TCDD in 5-month females for Hepatocytes, Stellate, Kupffer, NK-NKT, and Endothelial cells. GSEA core genes were further analyzed using ORA for enriched gene sets from the GOBP compendium. e. MuSiC analysis for cell composition of T2C-exposed livers. f. Heatmaps for overlap of core genes in attenuated aging in hepatocyte or nonparenchymal cell identity changes in exposed males (BPA.hi, TBT, and TCDD) or females (PM2.5-Chi, TBT, and TCDD).
