## Supplemental_Materials_Methods for "Early-Life Environmental Exposures Reprogram Epigenomic Aging to Alter Gene Expression Trajectories"

### SUPPLEMENTAL MATERIALS AND METHODS

#### Animal Exposures

BPA high dose, BPA low dose, DEHP, Pb, TBT, and TCDD and their corresponding diets and water were provided *ad libitum.* For BPA the modified AIN-93G diet supplemented with 50 mg/kg diet of BPA (diet TD.06156; Harlan Teklad) was used as or "high dose" and the modified AIN-93G diet supplemented with 50 µg/kg diet of BPA (diet TD.110337; Harlan Teklad) was used as the "low dose," as previously described [1]. Teklad Diets (Harlan Laboratories Inc, Madison, WI) provided all ingredients except BPA (Sigma Aldrich, St. Louis, MO). Drinking water was provided in polypropylene bottles. The estimated daily treatment of BPA in the study was 10 mg/kg body weight (bw) in the high dose and 10 µg/kg bw in the low dose groups.

DEHP was dissolved in corn oil from Envigo to create a customized stock solution, to produce 7% corn oil chow for experimentation. The DEHP exposure level was selected based on a target maternal dose of 5 mg/kg-day and assumes that a pregnant and nursing female mouse weighs approximately 25 g and ingests roughly 5 g of chow per day. This target dose was selected as previous literature demonstrated obesity-related phenotypes in offspring exposed to 5 mg/kg-day DEHP during early development [2], and this dosage falls within the range of exposures previously documented in humans [3]. Animals were maintained on phytoestrogen-free modified AIN-93 chow (Envigo Td.95092, 7% corn oil diet, Harlan Teklad) for the duration of the experiment.

Pb-acetate drinking water was prepared with distilled drinking water, with a concentration of 32 ppm to model human-relevant perinatal exposure. In previous work, we identified that this dose generates maternal blood levels (BLLs) ranging from 16-60 µg/dL (mean: 32.1 µg/dL) [4]. Exposure water was made by dissolving Pb (II) acetate trihydrate (Sigma-Aldrich) in a single batch of distilled water, and Pb concentrations were verified using inductively coupled plasma mass spectrometry with a limit of detection of 1.0 µg/L (ICPMS; NSF International). Animals were maintained on phytoestrogen-free modified AIN-93 chow (Envigo Td.95092, 7% corn oil diet, Harlan Teklad) for the duration of the experiment.

For TBT, animals were exposed to TBT chloride (TBT 96% purity; Sigma) in their drinking water, as previously described [5]. Consumption surveys were conducted in nonpregnant females to calculate the administered dose of TBT. Consumption levels were found to spike immediately after birth at the onset of lactation for a period of 3 d. Females were found to drink 10mL of water per day; therefore, they were given 200mL of 3.07µM TBT each week (equivalent to 0.5mg/kg BW per day) in the drinking water provided by the animal facility in typical plastic water bottles provided by the animal facility. This dose is below the lowest dose in the current developmental no observed adverse effect level (NOAEL) range of 5.8–20mg/kg BW as per the Concise International Chemical Assessment Documents (CICADs) guidelines for developmental toxicity [6]. Animals were maintained on phytoestrogen-free modified AIN-93G formulation (TD.94045) that replaces soybean oil with corn oil (Teklad) and fed *ad libitum* for the duration of the experiment.

TCDD – Female mice were exposed to TCDD (Cambridge Isotopes, cat#ED-901-C) dissolved in corn oil and administered by oral gavage. Dams were exposed in three doses. Each dose was 0.34 μg/kg body weight, for a total exposure of 1μg/kg body weight during the period from preconception to weaning. We administered pure corn oil to control mice. The doses were administered (1) at two weeks prior to mating; (2) approximately embryonic day 7; and (3) approximately postnatal day 14. Animals were maintained on phytoestrogen-free modified AIN-93 chow (Envigo Td.95092, 7% corn oil diet, Harlan Teklad) for the duration of the experiment.

Mice were exposed to ultrafine particulate matter (PM2.5) at two consortium sites: Johns Hopkins University (PM2.5-JHU) or University of Chicago. (PM2.5-Chi).

PM2.5-JHU, the PM2.5 or filtered air (FA) exposure was provided to the animals in the chambers as follows: 5 days per week (Mon-Fri) from 9 am to 5 pm (8 h/day) daily for 7 weeks. These animals were acclimated in our housing facility for one week before the chamber exposure. Each group of mice (FA or PM2.5) were regularly switched between the two separate exposure chambers throughout 7-week exposure period. There were no differences in locomotor activities or mobility in any of the mice assigned across groups prior to or during their exposure.

For PM 2.5-Chi, dams were placed daily for 8 hours in the PM or control chamber and were mated with male mice overnight until pregnancy is confirmed [7]. Once the pregnancy was confirmed, dams were exposed to either PM or control filtered air 8 hours per day until pups are born. Dams and pups were then exposed to PM or filtered air for 8 hours per day until the pups were 3 weeks old. The $PM_{2.5}$ was concentrated from ambient air in Chicago in a chamber connected to Versatile Aerosol Concentration Enrichment System (VACES) as previously described [8]. The chemical composition of PM has been previously reported [8]. We exposed control mice to filtered air in an identical chamber connected to the VACES in which a Teflon filter was placed on the inlet valve to remove all particles.
